# Distinct roles of brain network flexibility in motor learning across age

**DOI:** 10.1101/2024.10.29.620762

**Authors:** Kazumasa Uehara, Makoto Hagihara, Keiichi Kitajo

## Abstract

Motor learning is a lifelong process, from infancy through old age. Acquiring new motor actions through repetitive practice requires adjusting motor output in response to sensory input and integrating them to facilitate learning. For this to occur, the central nervous system must flexibly predict and adapt to the dynamic interplay between sensory inputs and motor outputs. Although overall brain function changes with age, it remains unclear how flexible brain networks, reflecting moment-to-moment reconfiguration of large-scale brain networks, influence motor learning ability with aging. To address this, we designed a visuomotor learning task involving both younger and older adults and quantitatively assessed brain network flexibility, leveraging multichannel electroencephalography (EEG) in humans. We found age-group differences in motor learning properties, brain network flexibility, and their neural relationships. In younger adults, the learning aftereffect was associated with brain network flexibility during motor preparation in the learning task. However, this association was not observed in older adults. Together, our findings suggest that brain network flexibility during motor preparation was critical for acquiring and maintaining new motor actions in younger adults, but that this coupling was attenuated with aging.

## Introduction

More recently, many people live well beyond the age of 60 and maintain good health. In our current reality, various types of motor and cognitive learning occur throughout a lifetime, spanning from early infancy to old age. Notably, learning new motor skills, such as playing musical instruments, remains feasible even in older adulthood. The definition of skill emphasizes reducing variability in action outcomes, and this variability progressively decreases throughout the learning process. However, the roles of movement variability and neural flexibility in motor learning remain subjects of ongoing debate. Although movement variability (i.e., exploratory behavior) is thought to facilitate motor learning (Ranganathan and Newell, 2010; Uehara et al., 2019; Ranganathan et al., 2020), and greater motor exploration has been linked to better learning outcomes (Wu et al., 2014), neural relationships remain unclear. A previous human electroencephalogram (EEG) study demonstrated that overall brain network flexibility reflects key neural processes involved in skill formation and predicts musical performance skill in proficient brass players. Notably, brain network flexibility primarily contributed to feedback control, rather than feedforward control, during skilled musical performance (Uehara et al., 2023). This study also utilized the brain network flexibility, which is defined as a quantitative index of node flexibility, capturing the frequency with which nodes change their module assignments over a given period. Here, nodes represent brain regions, and a module represents a group of brain regions that are more strongly interconnected with each other. This finding sheds new light on the crucial role of brain network flexibility in sensorimotor integration and the regulation of consolidated motor function. Although this analytical approach, namely brain network flexibility, is highly mathematical and initially appeared to be disconnected from biological mechanisms, accumulating empirical evidence suggests that it reliably captures fundamental properties of brain function (Braun et al., 2015; Betzel et al., 2017; Reddy et al., 2018; Monteiro et al., 2019; Uehara et al., 2023). Building on this body of evidence, we argue that brain network flexibility, reflecting moment-to-moment reconfiguration of large-scale brain networks, represents a key neural feature underlying motor learning.

From the perspective of aging, considering the age-related alterations in the neural system, as is widely accepted, age-related neural changes result in a gradual decline in sensorimotor function and learning ability (Vercillo et al., 2017; Hehl et al., 2020; Pauelsen et al., 2020). This decline is associated with reductions in white and gray matter volumes (Courchesne et al., 2000; Good et al., 2001; Jernigan et al., 2001) and alterations in white matter microstructural organization (Cox et al., 2016; Beck et al., 2021), changes in brain activity, and network dynamics (Garrett et al., 2013; Monteiro et al., 2019; Frolov et al., 2020; Goelman et al., 2023), and declines in dopaminergic and gamma-aminobutyric acid neurotransmission (Kaasinen and Rinne, 2002; Guitart-Masip et al., 2016; Hermans et al., 2018) even in healthy aging. These neural factors may individually or collectively disrupt the coordinated operation of brain networks across multiple levels, ultimately leading to behavioral changes (Marstaller et al., 2015; Shafiei et al., 2019; Yoshimura et al., 2020). There is substantial literature supporting theories of age-related changes in brain function, including findings of increased neural recruitment in the form of overactivation, increased connectivity, and frontal compensation. Of note, the Hemispheric Asymmetry Reduction in Older Adults (HAROLD) model, Posterior-Anterior Shift in Aging (PASA) theory, the Compensation-Related Utilization of Neural Circuits Hypothesis (CRUNCH), and the Scaffolding Theory of Aging and Cognition (STAC) serve as conceptual frameworks (Cabeza, 2002; Davis et al., 2008; Reuter-Lorenz and Cappell, 2008; Reuter-Lorenz and Park, 2014; Son et al., 2024).

Over the past decade, the concept of brain network flexibility has also emerged as an important framework for understanding age-related changes. For example, a human functional magnetic resonance imaging (fMRI) study has demonstrated that the strength of brain network flexibility was reduced in older adults, and the flexibility of the brain area in the prefrontal cortex was associated with motor skills including lower error during bimanual motor coordination (Monteiro et al., 2019). These findings allow us to raise an open question about whether age-group changes in brain network flexibility could hinder motor learning.

Here, we aimed to investigate whether age-group changes in brain network flexibility affect motor learning ability in both young adults (YA) and older adults (OA). We utilized multichannel scalp electroencephalogram (EEG) recordings while participants from both YA and OA groups engaged in learning an error-based visuomotor adaptation task. In the present study, we hypothesized that entire brain network flexibility would increase with aging due to excessive neural noise (Garrett et al., 2010) and compensatory reorganization (Cabeza, 2002). This increased flexibility becomes functionally ineffective in supporting motor learning in older adults. This is because higher brain network flexibility in older adults may reflect compensatory recruitment of neural resources rather than efficient network reconfiguration. Consequently, age-related neural deterioration may weaken the link between flexibility and the stabilization of motor memories, leading to a decoupling between neural flexibility and motor learning. We hypothesized that frameworks such as CRUNCH and STAC may provide a more compelling explanation than alternative frameworks, as mentioned above. Viewed holistically, focusing on whole-brain dynamics in terms of brain network flexibility is essential for further elucidating neural mechanisms of motor learning ability in the context of aging.

Moreover, an additional aim of this study was to determine how brain network flexibility changes over time during motor adaptation and to clarify its functional role in motor learning. Previous EEG studies have suggested that the suppression of trial-by-trial neural variability, as indexed by event-related potentials, was closely associated with enhanced perceptual abilities (Arazi et al., 2017; Daniel and Dinstein, 2021). Based on these findings, we hypothesized that the reduction in brain network flexibility during the motor learning process contributes to the stabilization of motor learning in young adults. Finally, we postulated that early consolidation of motor adaptation via wakeful rest (i.e., offline learning) may explain how brain network flexibility underpins a newly formed internal model of motor adaptation (Bönstrup et al., 2019; Buch et al., 2021). Using a short-term retention paradigm with a 10-min post-adaptation rest interval, we tested the hypothesis that brain network flexibility contributes to the formation and early stabilization of internal models during motor adaptation in both young and older adults.

## Materials and Methods

We utilized an error-based visuomotor adaptation task to assess individual motor learning abilities. To estimate the number of participants required to reliably assess motor learning effects, we conducted an a priori power analysis using the G*Power program (Erdfelder et al., 2009) with the effect size set at 0.49 based on our prior data collection from seven healthy right-handed young individuals (26.4 ± 5.5 yrs, range: 22 -35 yrs, two females). Using the same experimental protocol described later (see *Experimental task and protocols*), the effect size was determined assuming changes in motor performance (i.e., the degree of the deviation of cursor position) between the early, middle, and late phases of adaptation. The statistical significance level was set at p < 0.05 (two-tailed) for a one-way repeated measures analysis of variance (ANOVA) with a within-subject factor (adaptation phase). Despite the sample size calculation being conducted only for the younger group, this a priori power analysis estimation showed that at least 25 participants were required for one group, and effect sizes were reported for each statistic to confirm the robustness of the results. Based on this procedure, fifty-four healthy participants were recruited and divided into two groups, each consisting of 27 participants. The groups were categorized as younger-aged (YA) (27.6 ± 5.8 yrs, range: 20 - 38 yrs, 13 females) and older-aged (OA) (66.8 ± 7.4 yrs, range: 58 - 80 years old, 12 females). All participants were naïve concerning our visuomotor adaptation task, had a normal or corrected-to-normal vision, no history of musculoskeletal or neurological disorders, and were right-handed according to the Edinburgh Handedness Inventory (Oldfield, 1971). Before data collection, all participants in the OA group underwent a Mini-Mental State Examination (MMSE) with a cutoff score of 25 points, as scores above 25 points indicate intact cognition (Folstein et al., 1975). All participants in the OA group scored above 25 points (29.1 ± 1.2 points), leading to older adults having normal cognitive functions, including orientation, memory, attention, language, and visuospatial skills. However, four out of 27 participants from each group were excluded from the final data analyses due to the failure of the motor performance or electrical noise in the EEG data that was difficult to remove. This paper therefore reports data from 23 participants in each of the groups. This experimental protocol was approved by the local ethics committee of the National Institute for Physiological Sciences in accordance with the guidelines established in the Declaration of Helsinki. Before data collection, written informed consent was obtained from all participants.

### Experimental task and protocols

Experimental protocol, trial structure, and apparatus are illustrated in Figure 1. The electromagnetically shielded experimental room used in this study was completely darkened to minimize the availability of any visual information that was not projected on a PC monitor. Participants were seated 120 cm in front of a 24.5-inch (1920 x 1080 pixels resolution) computer display comfortably positioned on a chair with the right forearm supported by an armrest and a chin resting comfortably on a chin rest to ensure a consistent position throughout the experimental session.

**Figure 1:**
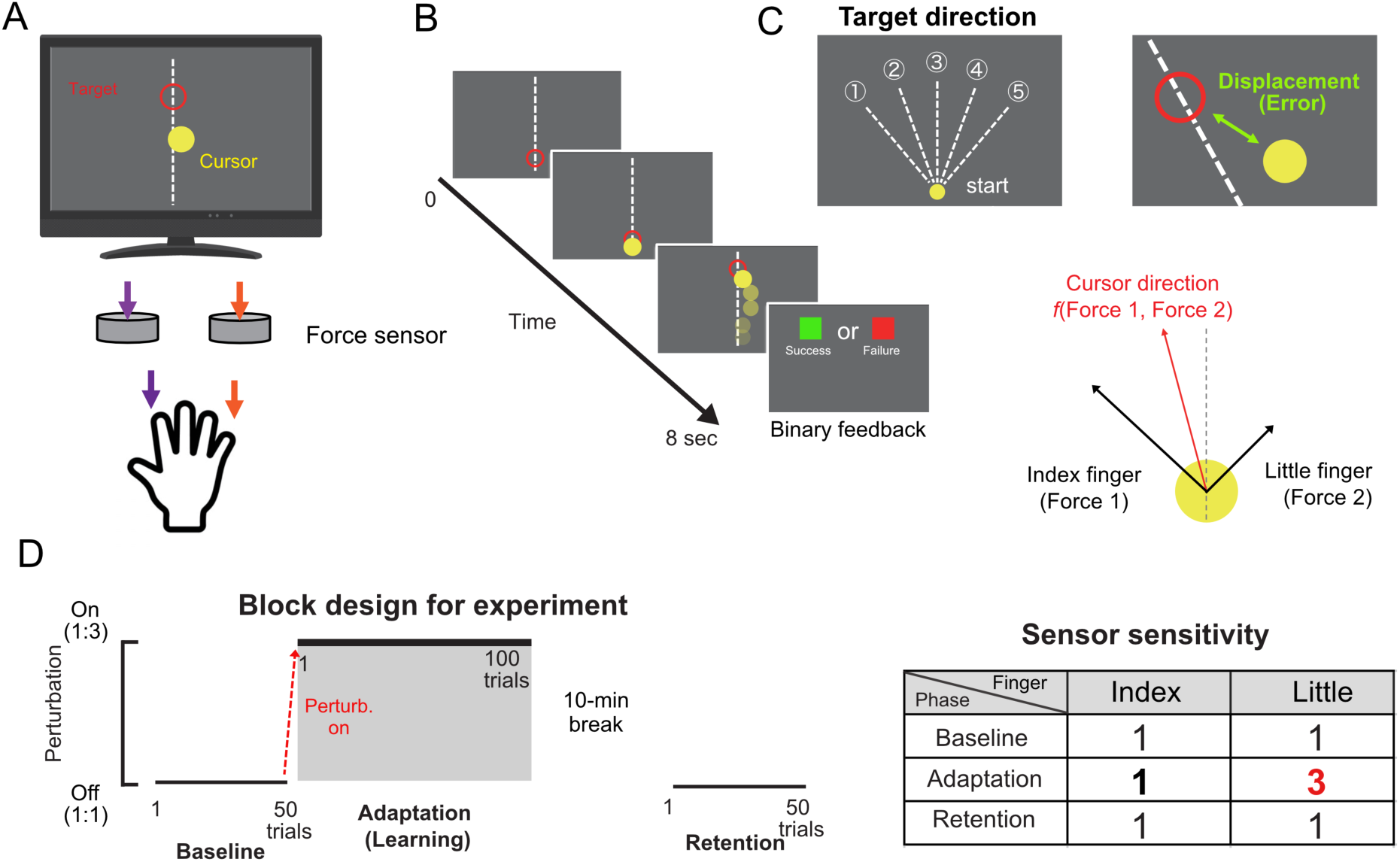
Experimental setup and trial structure. **A.** Participants were instructed to control a yellow cursor by pressing force transducers with their right index and little fingertips. **B.** Upon the ready cue (when the red open circle appeared), the yellow cursor was displayed on a monitor as the Go cue. After receiving the Go cue, participants began tracking the yellow cursor to keep the red circle on the target throughout the trial. **C.** The cursor direction controlled by a participant was determined using a composite vector related to the force exerted in the z-direction output from the index and little fingertips. At the end of each trial, they received binary feedback on their performance, with green or red squares indicating success or failure, respectively. The two-dimensional space distance (i.e., horizontal and vertical) between the moving target and the cursor was used as an error index. We employed an error-based visuomotor adaptation task with five different target directions (10, 11, 12, 1, or 2 o’clock direction). **D.** The sequence of all trials. In the first 50 trials, participants controlled the cursor with a force sensor sensitivity ratio of 1:1 for the index and little fingers during the baseline session. In the next 100 trials, which corresponded to the adaptation session, the force sensor sensitivity ratio was changed from 1:1 to 1:3 for the index and little fingers. After the 100 trials were completed, participants took a 10-minute break. Lastly, participants completed aftereffect testing consisting of 50 trials during which the force sensor sensitivity ratio was returned to the baseline session value (i.e., perturbation removal).

Prior to all data collection, participants underwent a 3-min resting-state EEG recording during which they remained at rest with their eyes open and were instructed not to focus on any particular thoughts. During the resting-state data collection, participants were instructed to see the fixation cross placed at the center of the monitor. Following this, participants performed three sustained maximal voluntary contractions (MVCs) of the right index and little fingers, respectively. The maximum force was defined as an MVC value and used to determine cursor control during the task.

On each trial, the target moved straight ahead at a constant speed of 10.8 cm/s in one of five directions (10, 11, 12, 1, or 2 o’clock) in a randomized order within each session. Participants were instructed to track a moving target on the screen by controlling the cursor as accurately as possible. Our experimental system allowed participants to freely control cursor speed and direction by pressing two force-torque transducers (USL06-H5, Tec Gihan, Co. Ltd., Japan) using their right index and little fingertips (Figure 1A). Our task design was inspired by previous literature (O’Sullivan et al., 2009), which demonstrated that the distribution of coordinated force made by the index and little fingers is greater when heteronymous effectors (e.g., index and little fingers) are used together than when homonymous combinations are used. This finding implies that the nervous system must adaptively coordinate the muscles by integrating both effort and variability. By employing a non-habitual effector like the little finger, we investigated how the brain flexibly learns force mapping in unfamiliar coordination contexts. The cursor direction controlled by a participant was determined using a composite vector related to the force exerted in the z-direction output from the index and little fingertips. In other words, by modulating the forces exerted by the index finger and little finger, participants could freely move the cursor in the left-right direction. Related to this, the cursor velocity was modulated according to the force applied in the z-direction of the force sensors, such that greater force resulted in faster cursor movement. Force signals were amplified via an amplifier device (GDA-06B, Tec Gihan, Co., Ltd., Japan) and then stored on a computer via a NI-DAQ device (USB-6002, National Instruments, United States) at a sampling frequency of 1 kHz with analog-to-digital conversion. The index finger controlled leftward cursor movement, while the little finger was responsible for rightward cursor movement. Therefore, the cursor direction was determined by the vector component calculated from the force applied by the index and little fingers. Cursor velocity was also controlled by varying the pressure applied to the force-torque transducers.

Visual cues, cursor control, cursor and target positions on the screen, and trigger signals sent to the EEG amplifier were managed by a dedicated software package (Graphical Design Lab, Japan), implemented in National Instruments LabVIEW software (version 2019, National Instruments, United States).

For the experimental protocol, participants were asked to learn the force mapping in their right index and little fingers through an error-based visuomotor adaptation task. The task began with a familiarization period consisting of 25 null movement trials performed without any added sensor perturbation. After the familiarization, we asked participants to complete a total of 200 trials. The experiment consisted of three conditions: baseline (unperturbed), adaptation (perturbed), and aftereffect. As shown in Figure 1B, at the beginning of each trial, the target circle appeared after a 5-second preparation period, signaling the “Go Cue”. Participants were asked to accurately track the target by controlling the cursor. At the end of each trial, they received binary feedback on their performance, with green or red squares indicating success or failure, respectively. Figure 1D illustrates the sequence of all trials. In the baseline condition consisting of 50 trials, participants tracked the moving cursor with a force sensor sensitivity ratio of 1:1 between the index and little fingers. During the adaptation phase, participants completed 100 trials with the force sensor sensitivity ratio of 1:3 between the index and little fingers, requiring them to adapt to the de novo force mapping. To investigate the relationship between brain network flexibility and early consolidation of motor adaptation induced by neural replay, we asked participants to complete an additional session without the sensor sensitivity changes after a 10-minute break. Previous motor adaptation studies have commonly employed retention tests after delays of several minutes (e.g., 10 min) to evaluate the stability of newly acquired motor memories (Bönstrup et al., 2019). From a molecular perspective, the 10-min rest, i.e., wakeful rest, likely captured the earliest stage of offline memory processing, during which consolidation-related molecular cascades may have already been initiated (Dudai, 2004; Parra et al., 2026). To do this, the aftereffect was assessed in a 50-trial session with sensor sensitivity returned to baseline after the 10-minute break. This additional session was included to evaluate whether an internal model had been sufficiently formed through motor adaptation, as the presence of the aftereffect is widely regarded as strong evidence of internal model formation. It is thought that the sensorimotor system adapts to the perturbation and updates its predictive control mechanisms, including an internal model (Salomonczyk et al., 2013; Leow et al., 2018; Krakauer et al., 2019; Tsay et al., 2024). Note that the luminance of the colors used for the cursor, target, and binary feedback was uniformly set to avoid any biases related to visual perception.

### Data recording

#### Behavioral data

Time series of the two-dimensional target and cursor positions (x- and y-axes) on the screen were continuously acquired at a sampling rate of 100 Hz using the dedicated software package as described above.

#### Electroencephalography

To record neural activity during data collection, we utilized a multi-channel EEG amplifier system (actiCHamp, Brain Products GmbH, Germany). EEG signals were continuously recorded from 63 electrodes positioned on the scalp following the international 10/10 system, using active electrodes embedded in a wearable elastic cap (actiCAP, Brain Products GmbH, Germany). A ground reference electrode for data acquisition was placed at AFz. During the data recording, all electrodes were referenced to the right earlobe, while the left earlobe was also recorded as an electrode for re-referencing. Electrooculography (EOG) signals are widely used to detect activities of eye movements and are available to remove eye movement-related noises from the EEG signals. EOG signals were recorded using electrodes placed above and below the left eye for monitoring vertical eye movements and below the left and right eyes for monitoring horizontal eye movements. EEG and EOG signals were amplified and digitized with 24-bit resolution at a sampling rate of 1 kHz. These signals were continuously stored on a personal computer via BrainVision recorder software (Brain Products GmbH, Germany). Skin/electrode impedance was consistently maintained below 10 kΩ throughout the data collection.

### Data analysis

Behavioral data were analyzed using custom-written code on MATLAB 2019b (MathWorks, Natick, United States). EEG data were also analyzed using functions from EEGLAB toolbox (Delorme and Makeig, 2004) implemented on MATLAB in combination with custom-written code. Statistical analyses for behavioral and EEG results were implemented using R environment (R Development Core Team, version 4.2.2, www.r-project.org/) unless otherwise noted.

#### Behavioral data

A two-dimensional space distance (i.e., screen height and width) between the moving target (red circle) and the cursor (yellow circle) was utilized as an error index. Throughout the error-based visuomotor adaptation task, participants had to reduce this error. To quantify the magnitude of the error, the Euclidean distance (*d*) from the moving cursor to the moving target was calculated at each timepoint (*t*) between the “Go” cue (*t*_0_) and end of the target tracking (*t*_end_) at each trial. We then calculated the cumulative error for each trial as follows:

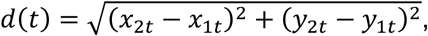

where *x*_1*t*_ and *y*_1*t*_ are the cursor center coordinates and *x*_2*t*_ and *y*_2*t*_ are the target center coordinates. Subsequently, the cumulative error (CE) for motor performance within each trial was calculated in the following formula:

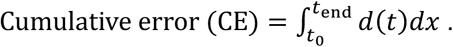

To examine the effects of motor adaptation, we computed the average error over five consecutive trials, defining this as one epoch.

With regard to quantifying motor adaptation, the CE data obtained during the adaptation period were first normalized to the average CE from the baseline period, i.e., null movement trials. This correction was performed because significant individual differences in basic motor skills among participants were observed, as shown in Supplementary Figure 1. By normalizing the data using the average of the baseline session, we minimized the influence of these differences in essential motor skills between participants, thereby facilitating a more accurate assessment of the adaptation process itself. Following this mathematical adjustment, the learning curve of errors across the trials in the perturbation phase was then fitted for each participant using an exponential function (Bönstrup et al., 2020) as follows:

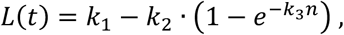

where *L* is the CE and *k*_1–3_ denotes initial performance (function intercept), maximum performance (function plateau), and learning rate (function slope), respectively. *n* represents the trial number.

To further determine how brain network flexibility relates to the ability to maintain and recall a learned motor performance, we computed the degree of aftereffect by averaging the error metric (i.e., displacement between the target and cursor) across the first five trials immediately after the 10-minute break. This is because the aftereffect is a hallmark of the effectiveness of motor learning. A high aftereffect value indicates that the motor performance and a related internal model have been solidly learned and can be recalled effectively.

#### EEG data

For preprocessing and artifact removal, consecutive EEG signals were re-referenced offline to the averaged signals of bilateral earlobes and segmented into 14-second epochs from 5 s before to 9 s after the “Go” cue. These segmented EEG data underwent band-pass filtering between 1-50 Hz while eliminating 60 Hz power line noise. Independent component analysis (ICA) was then applied to the filtered EEG signals associated with eye movements, blinks, and muscle contractions. The ICA-derived noise components were automatically identified using the ADJUST algorithm (Mognon et al., 2011). Across the individuals, at least two and up to six ICA components were removed from the filtered EEG signals (mean ± SD: 4.5 ± 1.1 components across participants). As a final preprocessing step, we applied the current source density transformation via the current source density toolbox (version 1.1), utilizing the spherical spline algorithm (Perrin et al., 1989; Kayser and Tenke, 2006). This step minimizes the impact of volume conduction on EEG data. Subsequently, artifact-corrected EEG data were used for brain oscillation analysis. Given the necessity of maintaining the sequence of motor adaptation processes, no trials were excluded to preserve the natural progression of motor adaptation.

To quantify functional brain networks and their network flexibility, we used the preprocessed EEG data obtained from 63 channels and computed phase-based measures of neural oscillatory activity (Lachaux et al., 1999; Varela et al., 2001; van Diessen et al., 2015). This approach aligns with the dynamical systems observed in the brain (Lachaux et al., 1999), where phase synchrony of neural activity significantly contributes to information processing within the brain (Rodriguez et al., 1999; Varela et al., 2001; van de Vijver and Cohen, 2019). First, to decompose the single-trial EEG signals into time-frequency representations of the instantaneous phase, we applied complex Morlet wavelet that can be defined as the product of a complex sine wave and Gaussian window (Tallon-Baudry et al., 1996; Cohen, 2019). The CSD-transformed EEG signals were convoluted by a Morlet wavelet function *w*(*τ*, *f*):

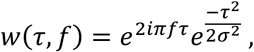

where *i* is the imaginary unit. *τ* and *f* denote time points and the center frequency, respectively. *σ* is the width of the Gaussian, which is defined as

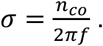

The parameter *n_co_* is defined as the number of cycles and represents a nontrivial choice in time–frequency analysis. In the present study, we set *n*_co_ = 3 and *f* increased from 1 to 50Hz in 1-Hz steps (Lachaux et al., 2000). It is well established that this parameter governs the trade-off between temporal and frequency precisions, such as that a larger number of cycles results in lower temporal resolution, and vice versa. A smaller number of *n*_co_ enables more precise localization of temporal dynamics compared to a larger number. Our primary aim is to capture transient changes in neural activity within a time-limited window during motor execution associated with learning. Drawing on established knowledge (Cohen, 2014), we therefore selected a smaller number of cycles. We then calculated the instantaneous phase synchrony index (PSI) at each frequency for all electrode pairs using the following formulas;

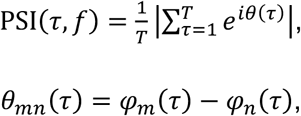

where *T* denotes the analysis window length, which was fixed at 500 ms (i.e., 500 data points) for all frequency bands, using 90% overlapping windows, following previous literature (Uehara et al., 2023; Krukow et al., 2024). This window length was selected to include at least three oscillatory cycles of the slowest frequency (e.g., Theta frequency band), thereby striking a balance between reliable phase synchronization estimation and the temporal resolution required to capture dynamic brain network changes (Liuzzi et al., 2019). In addition to this, the utilization of 90% overlapping windows enables the generation of time-dense phase synchrony information, facilitating the observation of highly time-resolved brain network dynamics during the task. *φ_m_*(*τ*) and *φ_n_*(*τ*) represent the instantaneous phase of the *m_t_*_ℎ_ and *n_t_*_ℎ_ electrodes at time point *τ*. The PSI represents phase synchronization between EEG signals from two different electrodes at each time point within each time window. This index ranges between 0 and 1, where a value of 0 indicates complete randomness, while a value of 1 denotes perfect synchrony (i.e., complete phase alignment) in EEG signals from the two electrodes and provides valuable information about functional connectivity within the whole brain (Lachaux et al., 2000; Varela et al., 2001).

As the final step, the whole-brain network flexibility was determined using the Louvain-like algorithm (Blondel et al., 2008; Mucha et al., 2010), which is a mathematical method for revealing the inherent module structure within complex networks. This approach is rooted in network analysis, employing graph theory and has recently been applied to analyze neuroimaging data, including fMRI and EEG, which is unveiling functional relationships between brain network flexibility and cognition as well as action (Bassett et al., 2011, 2013; Betzel et al., 2017; Paban et al., 2019; Uehara et al., 2023). Initially, the generalized Louvain algorithm was employed for multilayer module detection. In the present study, a multilayer module was defined wherein each network node (i.e., EEG electrode) in one time window is connected to itself in neighboring (preceding or following) time windows (i.e., layers). This step can be calculated using the following formula (Mucha et al., 2010; Bassett et al., 2013):

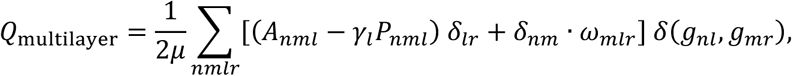

where *Q* represents the multilayer modularity index, *l* denotes the number of layers in the multilayer network, and *μ* signifies the total edge weights encompassing both intra- and inter-layer connections. *A_nml_* represents the time-by-time *N* × *N* PSI-derived functional connectivity matrices (where *N* denotes the number of EEG electrodes) between nodes *n* and *m* (i.e., PSI value). *P_nml_* signifies the weight of the edge between nodes *n* and *m* under the null model, while *γ_l_* denotes the structural resolution parameter that defines the weight of the intralayer connections. *g_nl_* represents the module assignment of node *n* in layer *l* while *g_mr_* is the module assignment of node *m* in layer *r*. *ω_mlr_* denotes the connection strength between nodes across two layers. In this study, the parameters *ω* and *γ* were set to 1, constituting their default values as per previous literature (Bassett et al., 2013). *δ*(*g_nl_*, *g_mr_*) corresponds to the Kronecker delta, taking a value of 1 if *g_nl_* = *g_mr_* and a value of 0 if *g_nl_* ≠ *g_mr_*. Given the heuristic nature of this algorithm, we ran the analysis 100 times and selected the best clustering based on the maximum Q value. Subsequently, we determined brain network flexibility (*f_n_*) by counting how often a node altered its module assignment throughout time, adjusting for the total potential changes (*N*) by using the following formula,

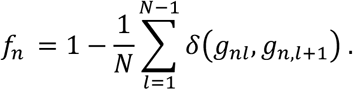

Brain network flexibility (*f_n_*) was averaged in the theta (4-8 Hz), alpha (9-13 Hz), and beta (14-30 Hz) frequency bands across all EEG electrodes to capture the global flexibility of the whole cortical regions on every single trial. This flexibility index ranges from 0 to 1, where 0 indicates no change in communities, while 1 indicates a change in the module across every layer. Before averaging across trials, the flexibility of the whole brain network was averaged within each trial across predefined frequency bands during the 5-second motor preparation period immediately preceding the onset of action and during the 3-second motor execution period. We presumed that the neural systems dynamically and flexibly reconfigure brain networks not only during motor execution but also during the preparation period throughout motor adaptation. Previous human behavioral studies demonstrated that the preparation of motor execution during a motor adaptation process plays a critical role in the ability to learn in an adaptation task (Haith et al., 2015). Preparatory neural activity patterns have the potential to retain motor memory (Sun et al., 2022). Focusing more directly on brain network flexibility, preparatory brain network flexibility for musical playing is necessary for feedback control, rather than feedforward control, in skilled motor performance by expert musicians (Uehara et al., 2023). These previous studies motivated us to investigate how the flexibility of neural preparatory activity affects motor learning ability across different ages. Moreover, we quantified temporal changes in brain network flexibility throughout the trials during adaptation to identify how inter-trial brain network flexibility changes within the adaptation period and how these changes relate to motor learning ability with age. We performed this analysis to test the assumption that reduced flexibility may stabilize brain networks optimized by learning and enhance focus on relevant neural signals. This hypothesis is supported by a previous human EEG study, which suggests that the suppression of trial-by-trial neural variability is closely associated with perceptual abilities (Arazi et al., 2017). To test this, we calculated the differences in brain network flexibility values between the averages of the first and last 5 trials according to the predefined frequency bands at the individual level. Additionally, the resting-state EEG data were also analyzed by using the same procedures as those used for the task-related EEG analysis described above.

To enhance the interpretability of brain network flexibility in greater depth, we computed the multi-channel Lempel-Ziv complexity (LZC) index (Lempel and Ziv, 1976; Fernández et al., 2011). This index quantifies the dynamical complexity generated by brain networks in terms of randomness and compressibility. There is a possibility that our mathematical approach merely captures the apparent flexibility of brain networks. In other words, even if network nodes transition frequently within a small number of modules, our approach might incorrectly determine that the brain network is highly flexible. We assessed LZC in each frequency band across the group to rule out this false detection. Multi-channel LZC was computed for motor preparation and motor execution periods of each trial using a code written in MATLAB. We then calculated the trial-averaged LZC at the individual level. LZC during the resting state was also computed in the same manner. In general, higher LZC values (normalized between 0 and 1) reflect greater randomness. Additionally, although the present study has a potential limitation in topographical interpretation since we did not perform source localization analysis, looking at the topographical characteristics of each network module will help in biologically interpreting the obtained results. To do so, leveraging insights into the module affiliations of each node (i.e., EEG channel) over the analytic time window, we visualized each network module associated with brain network flexibility in each group using a hierarchical clustering algorithm (function *cluster* in MATLAB).

### Statistics

#### Motor learning profiles

To test age-group differences in the motor learning profiles between the YA and OA groups, we used an independent-sample t-test for the learning slope and aftereffect, respectively.

#### Brain network flexibility and LZC

To test age-group differences in brain network flexibility and LZC during the resting state, we separately conducted a two-way repeated measures ANOVA with a mixed design on brain network flexibility and LZC in the resting state with the AGE (2 levels: YA and OA) as the between subject-factor and FREQUENCY (3 levels: theta, alpha, and beta) as the within subject-factor. Next, a three-way repeated-measures ANOVA with a mixed design was conducted on brain network flexibility and LZC obtained during motor adaptation, with AGE (2 levels: YA and OA) as the between-subject factor and STATE (2 levels: motor preparation and motor execution) and FREQUENCY (3 levels: theta, alpha, and beta) as the within-subject factors. Moreover, we are interested in the extent to which changes in brain network flexibility are associated with the progression of motor learning. To test this, a four-way repeated measures ANOVA for mixed design with the AGE (2 levels: YA and OA) as the between subject-factor and STATE (2 levels: preparation and motor execution), FREQUENCY (3 levels: theta, alpha, and beta) and PHASE (2 levels: mean of the initial and final 5 trials during the perturbation session) as the within subject-factors was conducted. A Greenhouse-Geisser correction for a series of ANOVAs was used when the assumption of sphericity was violated. Post hoc explorations using the Holland-Copenhaver correction for multiple comparisons were performed if a significant effect was detected from the ANOVA (Holland and Di Ponzio Copenhaver, 1988).

Furthermore, we determined relationships between brain network flexibility and LZC by using Pearson correlation analyses. The correlation analyses between variables of brain network flexibility and LZC were conducted separately for each condition and the age groups. The reported p-values (< 0.05) were corrected for multiple comparisons using the Bonferroni correction.

#### Neuro-behavioral relationship

After characterizing condition-dependent variations in brain network flexibility and LZC, we next examined whether brain network flexibility during the resting state, motor preparation, and motor execution across frequency bands was associated with motor learning ability, as assessed by learning rate and aftereffect. In addition, we examined whether changes in brain network flexibility over the course of learning were related to motor learning performance. To identify the neural variables that most strongly contributed to motor learning outcomes, we performed least absolute shrinkage and selection operator (LASSO) regression analyses. LASSO regression enables the selection of explainable variables using a weighted combination of L1 norm regularization, which incorporates penalized regression (Tibshirani, 1996). The LASSO regression used in the present study was performed using the *glmnet* package implemented in R software package (Friedman et al., 2010). Separate LASSO models were constructed for the resting state, motor preparation, and motor execution. In each model, the trial-averaged brain network flexibility values at theta, alpha, and beta frequencies were treated as independent variables. The individuals’ learning rate and aftereffect were treated as dependent variables, respectively. To ensure the goodness-of-fit of the model, a coefficient of determination value (*R*^2^) was used. We repeated the regression 100 times and identified the median of R^2^ across these repetitions because the results of LASSO regression are indeed sensitive to the choice of the hyperparameter. These repetitions can help to provide a more robust estimate of model performance. The optimal *λ* was chosen by performing the default 10-fold cross-validation and selecting the one with the minimum generalization error. Based on this median *R*^2^ obtained from each LASSO model, we identified and reported the brain network flexibility measures most strongly associated with motor learning profiles.

## Results

### Behavioral results

Overall motor learning profiles are shown in Figure 2A. Representative cursor trajectories during motor adaptation are depicted in Supplementary Figure 1B. Figure 2A shows the degree of behavioral performance across motor adaptation observed in the YA and OA groups. Motor learning was evaluated by the degree of baseline-normalized performance error. This is then transformed into parameters representing each profile. As predicted, the degree of error was attenuated along with the number of trials, and this was visible from the first few trials after the start of the adaptation period, even though baseline accuracy (i.e., first 50 trials without the perturbation) differed between the groups (Supplementary Figure 1A). This indicates that motor learning occurred throughout the adaptation period in both groups. Subsequently, we found that the learning rate, parameterized by the slope (i.e., β value) of an exponential fit per individual, significantly differed between the groups (Figure 2B) according to an independent t-test (*t* = 2.574, *p* = 0.013, Cohen’s d = 0.75). The YA group exhibited a lower slope value compared to the OA group, indicating that the YA group was able to sustainably attenuate performance error throughout the adaptation period. No significant difference in aftereffect was detected between the groups (*t* = -0.504, *p* = 0.616, Cohen’s d = 0.42) (Figure 2C). These results suggest that age had minimal impact on the effectiveness of motor learning and the ability to recall motor memory.

**Figure 2:**
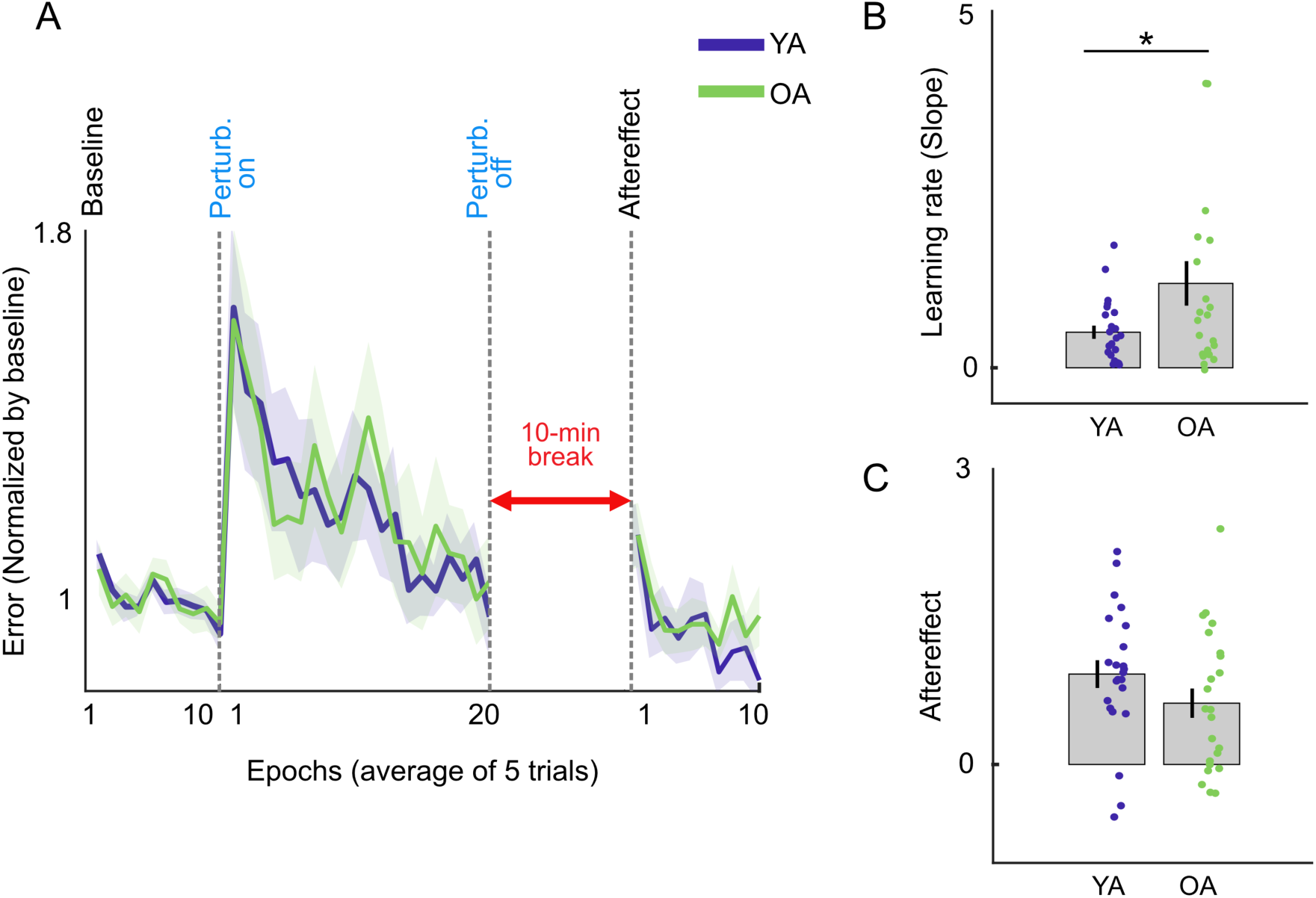
Behavioral results. **A.** Group-averaged performance error normalized by the baseline value throughout the data collection. Solid lines and shaded areas showed mean and SEM for five trials of epoch error (displacement between the target and cursor across participants (n=23 of each). **B.** Learning rate was determined by the slope value (*k*^3^) of the exponential function. Group-averaged learning rates significantly differed between the groups. A lower value indicates continued performance improvement throughout the adaptation period. **C.** Group-averaged aftereffect was used as a hallmark of the effectiveness of motor learning. We calculated the degree of aftereffect by averaging the first five trials after the 10-minute break. A higher value indicates that the motor performance has been solidly learned and can be recalled effectively. Each dot represents an individual, and each error bar indicates the standard error of the mean across participants. Statistical significance is denoted by ∗ p < 0.05.

### Brain network flexibility

We first sought to verify how brain network flexibility at the resting state differs between the two groups (Figure 3). A two-way ANOVA found a significant main effect of FREQUENCY (*F*_1.9,87.2_ = 10.53, p < 0.001, 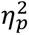 = 0.19), but no significant AGE effect (*F*_1,44_ = 0.82, p = 0.36, 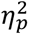 = 0.01). The significant interaction between them was detected (*F*_1.9,87.2_ = 3.52, *p* = 0.035, 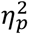 = 0.07). Post hoc tests revealed that the group differences in the alpha frequency band did not reach statistical significance (*F*_1,44_ = 3.95, p = 0.0529, 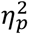 = 0.08). The theta frequency band showed significantly higher brain network flexibility than the other frequency bands in the YA group, but not in the OA group (*p* < 0.05, Holland-Copenhaver corrected).

**Figure 3:**
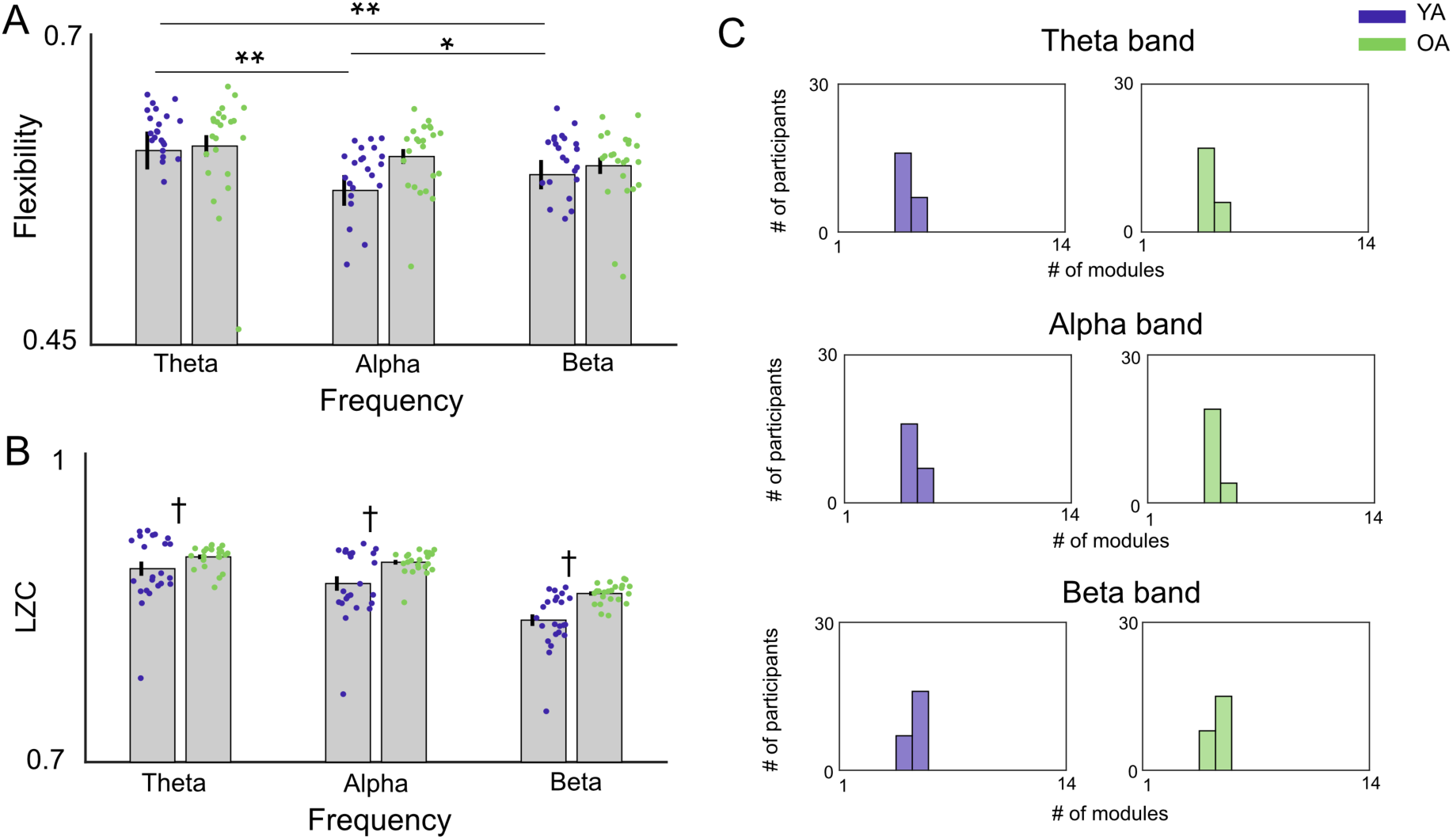
Group-averaged brain network flexibility during an eyes-open resting state in response to the predefined frequency bands **(A)** and their network properties (**B**: LZC, **C**: Number of network modules). Each dot represents an individual, and each error bar indicates the standard error of the mean across participants. Statistically significant differences between the frequency bands are denoted by ∗∗ p < 0.001 and ∗ p < 0.05. A statistically significant difference between the age groups is indicated by † p < 0.05.

Next, we tested whether age-group differences in brain network flexibility emerged during motor preparation (Figure 4A) and motor execution (Figure 4E) periods throughout the motor adaptation session. A three-way ANOVA found significant main effects of the STATE (*F*_1, 44_ = 11.11, p = 0.001, 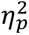 = 0.20) and FREQUENCY (*F*_1.48,64.9_ = 25.66, p < 0.001, 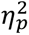^2^ = 0.36), but no main effect of AGE (*F*_1,44_ = 0.425, p = 0.42, 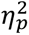 = 0.014). A significant interaction between STATE and FREQUENCY was detected (*F*_1.27,55.9_ = 50.97, p < 0.001, 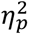^2^ = 0.53). No significant interaction effects involving the age factor were observed (all p > 0.05). Despite no age-group differences being observed, significant differences in brain network flexibility were observed between the preparation and motor execution periods (Theta: *F*_1,44_ = 33.06, p < 0.001, 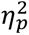 = 0.42; Alpha: *F*_1,44_ = 64.08, p < 0.001, 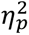 = 0.59; Beta: *F*_1,44_ = 15.74, p < 0.001, 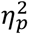 = 0.26). In the present study, another important aim was to examine how the degree of brain network flexibility changed over time throughout the adaptation period.

**Figure 4:**
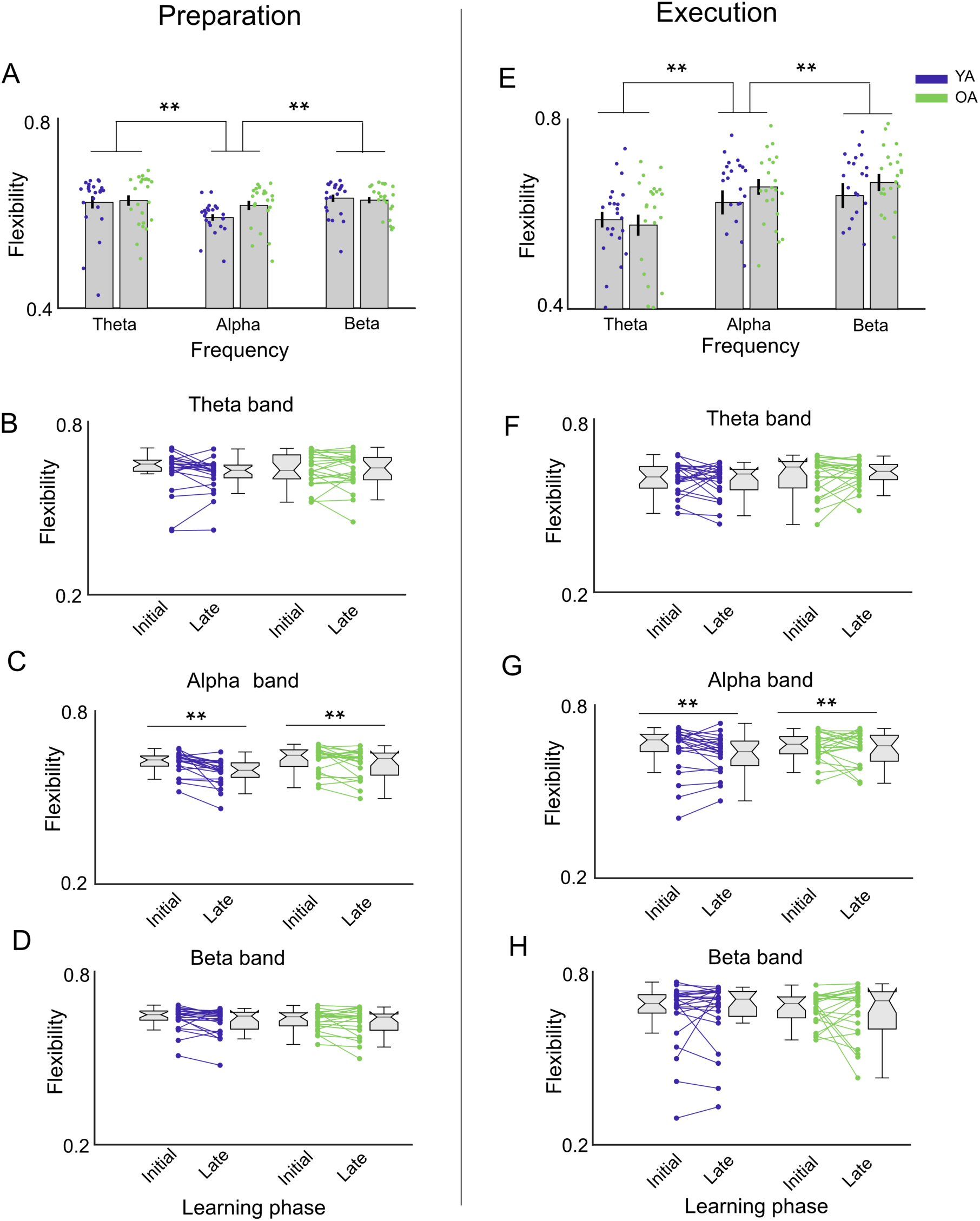
Group-averaged brain network flexibility during the preparatory period **(A)** and motor execution **(E)** in the visuomotor learning task in response to the predefined frequency bands. A higher value indicates greater flexibility among the networks. **B-D.** Changes in group-averaged brain network flexibility during the preparatory period from the initial to the late trials were compared for each frequency band (B: Theta, C: Alpha, and D: Beta). **E-H.** show similar patterns, but the results correspond to the motor execution (F: Theta, G: Alpha, and H: Beta). Each dot represents an individual, and each error bar indicates the standard error of the mean across participants. Statistical significance is denoted by ∗∗ p < 0.001.

Figure 4B-D and Figure 4F-H illustrate how brain network flexibility changes from the initial to the later phase over the time course of the adaptation period. The four-way ANOVA detected significant main effects of PHASE (*F*_1,44_ = 7.371, *p* = 0.009,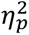^2^= 0.14) and FREQUENCY (*F*_1.45, 63.88_ = 20.93, *p* < 0.001, 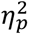= 0.32) but no significant main effects of STATE (*F*_1,44_ = 1.43, *p* = 0.237, 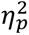 = 0.03) or AGE (*F_1_*_,44_ = 0.804, *p* = 0.374, 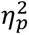 = 0.01). Moreover, a significant interaction between STATE and FREQUENCY was detected (*F*_1.62,71.48_ = 27.58, *p* < 0.001, 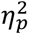= 0.385), whereas no other statistically significant interactions were detected (all *p* > 0.05). To further examine the significant interaction observed in the four-way ANOVA, follow-up post hoc tests were conducted. Brain network flexibility was significantly modulated depending on the initial or late phases of adaptation (*F*_1, 44_ = 13.22, *p* < 0.001, 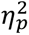 = 0.23). Brain network flexibility in the alpha frequency band significantly decreased toward the end of the adaptation period (*F*_1, 44_ = 23.46, *p* < 0.001, 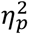 = 0.34). On the other hand, during motor execution, there was no significant difference in brain network flexibility between the initial and late phases of the adaptation period (*F*_1, 44_ = 0.63, *p* = 0.428, 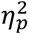 = 0.01). Taken together, these findings suggest that brain network flexibility decreased as learning progressed, indicating a gradual stabilization of network dynamics during the motor learning process.

To gain additional insights into the characteristics of the functional brain network, we assessed how many network modules were mediated by the flexible nodes because there was a concern that the network could be considered highly flexible even if each node was only moving back and forth between a limited number of modules. To address this concern, the number of modules and network complexity, as indicated by LZC, were reported, respectively. For the resting-state brain network, the two-way ANOVA for the LZC values detected significant main effects of AGE (*F*_1, 44_ = 9.77, *p* = 0.003, 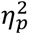= 0.18) and FREQUENCY (*F*_1.45,63.71_ = 619.2, *p* < 0.001, 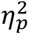= 0.93) and their significant interaction (*F*_1.45,63.71_ = 20.49, *p* < 0.001, 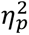= 0.31), as shown in Figure 3B. The OA group exhibited higher dynamical complexity, as reflected by brain network flexibility, than the YA group, suggesting that aging may be accompanied by increased randomness in resting-state brain network dynamics. Post hoc test detected significant age-group differences between the alpha and beta bands, but not the theta band (Theta: *F*_1, 44_ = 2.48, *p* = 0.122, 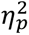= 0.05; Alpha: *F*_1, 44_ = 12.33, *p* = 0.001, 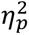= 0.21; Theta: *F*_1, 44_ = 18.67, *p* < 0.0001, 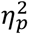= 0.29).

Figure 5B shows the LZC values during the motor preparation and motor execution periods. The three-way ANOVA for the LZC values detected significant main effects of STATE (*F*_1, 44_ = 174.3, *p* < 0.001, 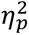= 0.79) and FREQUENCY (*F*_1.37,60.12_ = 221.6, *p* < 0.001, 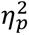 = 0.83) as well as significant interaction between them (*F*_1.53,67.11_ = 55.71, *p* < 0.001, 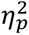= 0.55). More concretely, the LZC values in motor execution were significantly lower than those in motor preparation across all frequencies. In contrast, no significant effects or interactions involving the factor AGE were observed (all *p* > 0.05). Table 1 summarizes the correlations between LZC and brain network flexibility during the resting state, motor preparation, and motor execution in both groups. Significant positive correlations were observed across all conditions and frequency bands, except for the beta band in the OA group. These results suggest that local signal complexity, which corresponds to LZC, and large-scale network reconfiguration, which corresponds to brain network flexibility, may be constrained by a common neural dynamical process.

**Figure 5:**
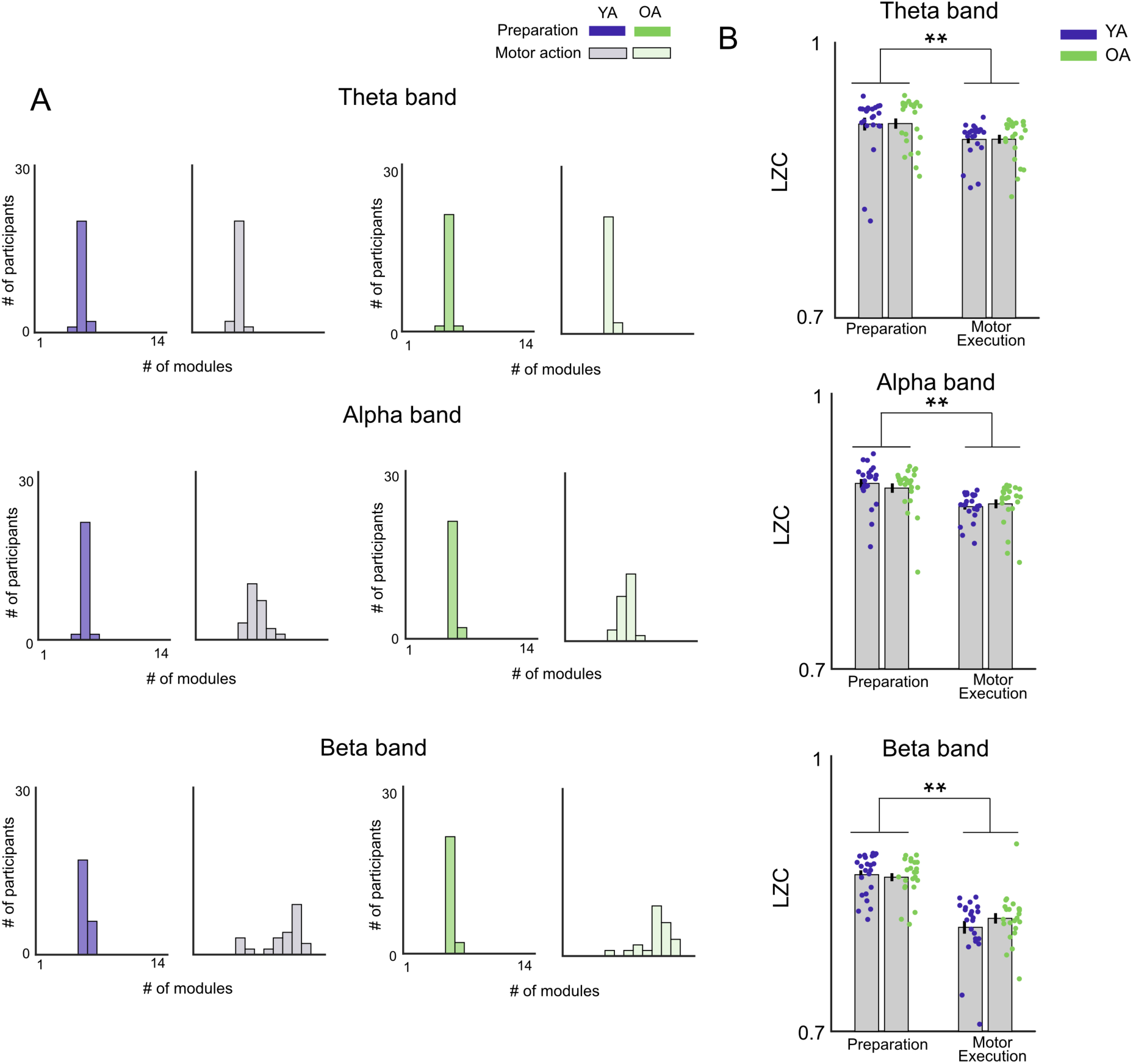
Characteristics of brain network structures. **A.** Number of modules in the extracted brain networks for predefined frequency bands under the preparatory and motor execution period during the motor learning. **B.** Comparisons of brain network complexity. A higher value indicates that the extracted brain networks are complex and contain many unique patterns. Each dot represents an individual, and each error bar indicates the standard error of the mean across participants. Statistical significance is denoted by ∗∗ p < 0.001.

**Table 1:** Correlations between LZC and Brain Network Flexibility.

|  | Resting state |  | Motor Preparation |  | Motor Execution |  |
| --- | --- | --- | --- | --- | --- | --- |
|  | YA | OA | YA | OA | YA | OA |
| Theta | $r = 0.76$<br>$p = 4.56 \times 10^{-4}$ | $r = 0.91$<br>$p = 2.32 \times 10^{-8}$ | $r = 0.85$<br>$p = 3.22 \times 10^{-6}$ | $r = 0.69$<br>$p = 3.7 \times 10^{-3}$ | $r = 0.85$<br>$p = 4.26 \times 10^{-4}$ | $r = 0.84$<br>$p = 0.001$ |
| Alpha | $r = 0.82$<br>$p = 2.50 \times 10^{-5}$ | $r = 0.73$<br>$p = 0.0012$ | $r = 0.59$<br>$p = 0.0494$ | $r = 0.76$<br>$p = 3.39 \times 10^{-4}$ | $r = 0.59$<br>$p = 0.015$ | $r = 0.72$<br>$p = 7.99 \times 10^{-6}$ |
| Beta | $r = 0.79$<br>$p = 1.10 \times 10^{-4}$ | $r = 0.60$<br>$p = 0.0424$ | $r = 0.96$<br>$p = 4.92 \times 10^{-12}$ | $r = 0.90$<br>$p = 6.54 \times 10^{-8}$ | $r = 0.96$<br>$p = 2.36 \times 10^{-6}$ | $r = 0.25$<br>$p = 4.34$ |

Note: Upper rows show correlation coefficients ( $r$ ), and lower rows show corresponding adjusted $p$ values.

### Neuro-behavioral relationship

Figures 6 and 7 summarize the relationships between brain network flexibility and the learning properties based on the results of the LASSO regression analyses. Figure 6A shows the extent to which brain network flexibility during the preparation period predicted the behavioral outcomes. We found that brain network flexibility across the entire frequencies accounted for 19.3%, [interquartile range (IQR): 0.5%] of the aftereffect in YA, whereas this relationship was scarcely explainable in OA (2.5%, [IQR: 0.1%]). Interestingly, the learning rate in OA was partially explained by brain network flexibility in the theta and beta bands (11.4%, [IQR: 0.1%]). Figure 6B shows the relationships between changes in brain network flexibility toward the end of the adaptation period and the behavioral outcomes. We found that the degree of change in brain network flexibility during the adaptation period predicted learning even more strongly. In the YA group, this relationship was highly explainable for a substantial portion of the aftereffect (51.2%, [IQR: 0%]), whereas in the OA group, it was primarily associated with the learning rate (25%, [IQR: 0.3%]). On the other hand, brain network flexibility during motor execution predicted only the learning rate, but not the aftereffect (Figure 7). Furthermore, its explanatory contribution of brain network flexibility during motor execution was clearly lower than during preparation across the age groups. For comparison, brain network flexibility during the resting state did not predict either learning rate (YA group: 7.0% [IQR: 0%], OA group: 0% [IQR: 0%]), or aftereffect (YA group: 4.2% [IQR: 0%], OA group: 8.5% [IQR: 5.2%]) in either age group. Importantly, the apparent discrepancy arises because the ANOVA and LASSO analyses address different questions. The ANOVA tested whether phase-dependent changes differed between age groups at the group level, whereas the LASSO analyses identified variables that best predicted individual differences within each age group. Therefore, age-specific predictive relationships may emerge even in the absence of a significant AGE and PHASE interaction. Our findings highlight that brain network flexibility during motor preparation contributed more to motor learning in younger adults than in older adults, whereas brain network flexibility during motor execution showed a weaker contribution.

**Figure 6:**
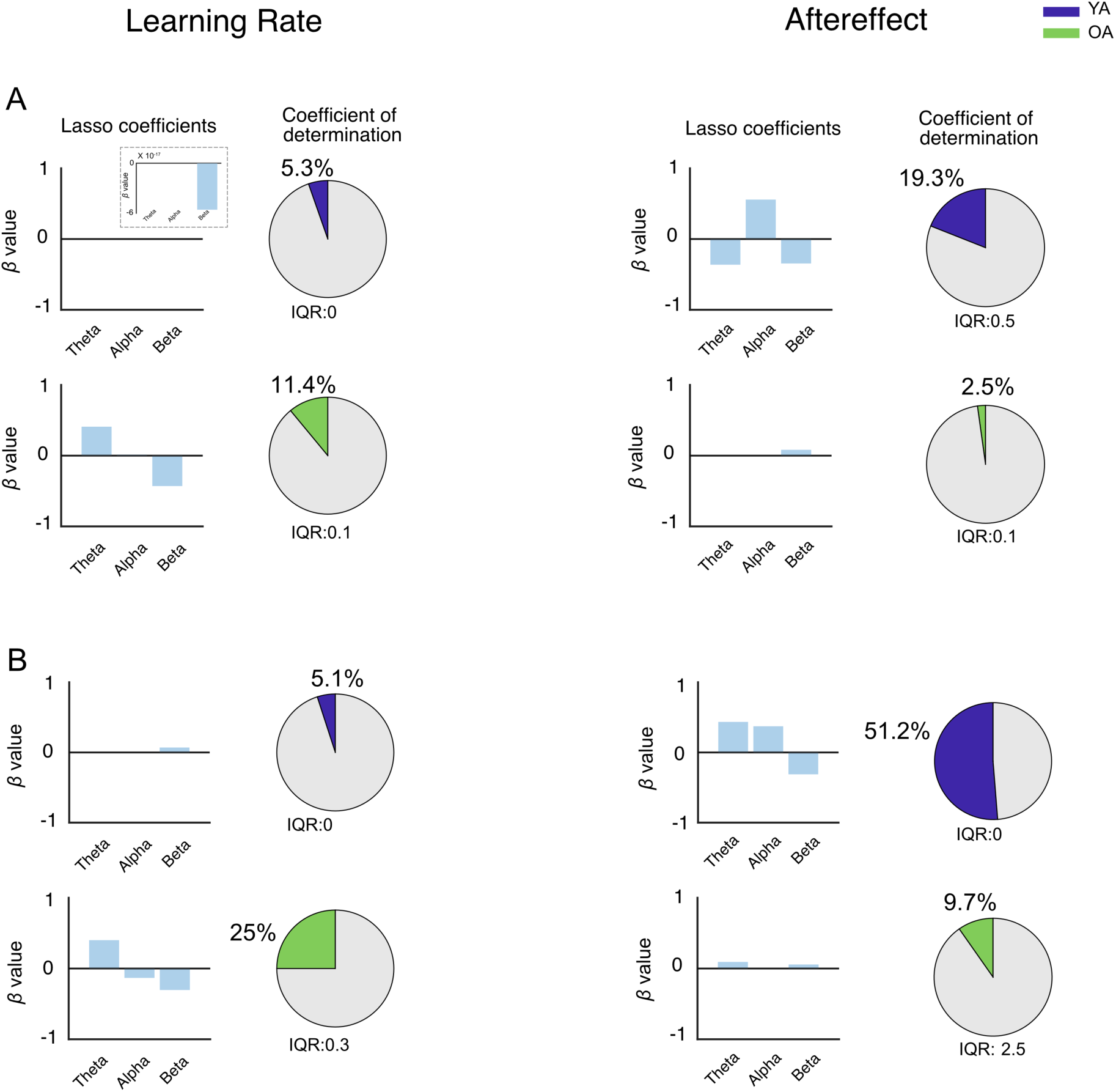
Results of the LASSO regression during the preparatory period. **A.** Relationships between brain network flexibility during the preparation period and motor learning outcomes. **B.** Relationships between temporal changes in brain network flexibility throughout the adaptation period and motor learning outcomes. Bar plots display β coefficients estimated by LASSO, while pie charts summarize the explanatory power as the coefficient of determination (R^2^). The small inset within the bar plot for the β coefficients provides an enlarged view of the region near zero. The interquartile range (IQR) of each R² value is shown at the bottom of the corresponding pie chart.

**Figure 7:**
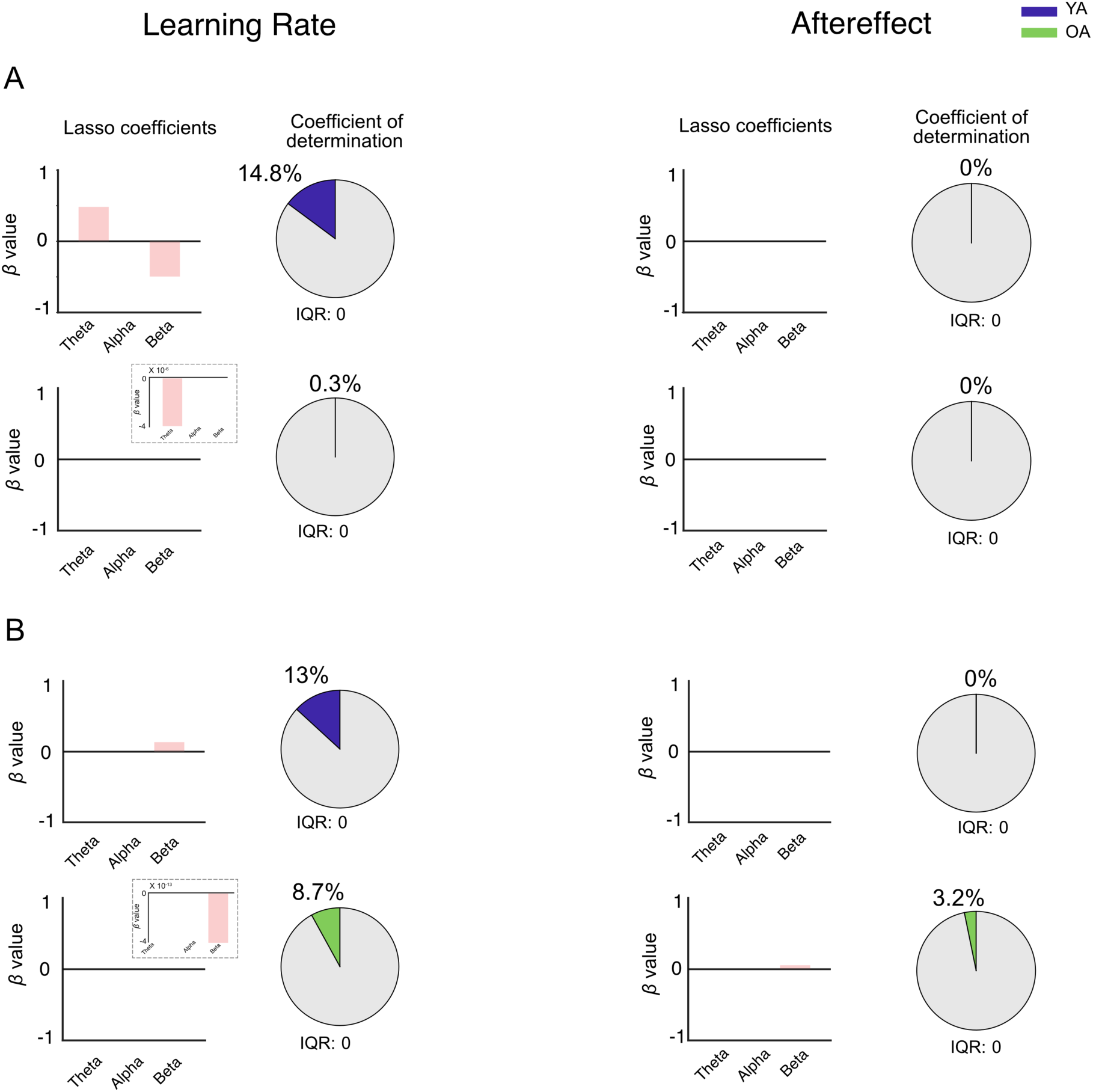
Results of the LASSO regression during the motor execution. **A.** Relationships between brain network flexibility during the motor execution and motor learning outcomes. **B.** Relationships between temporal changes in brain network flexibility throughout the adaptation period and motor learning outcomes. Bar plots display β coefficients estimated by LASSO, while pie charts summarize the explanatory power as the coefficient of determination (R^2^). The small inset within the bar plot for the β coefficients provides an enlarged view of the region near zero. The interquartile range (IQR) of each R² value is shown at the bottom of the corresponding pie chart.

## Discussion

The present study demonstrated that brain network flexibility was a key neural correlate of motor learning abilities, although the nature of this relationship differed between the age groups (Figures 6 and 7). First, brain network flexibility during motor preparation was more strongly associated with consolidation of motor learning than that during motor execution. Second, these associations were more pronounced in younger adults than in older adults. Third, a gradual decrease in brain network flexibility (i.e., stabilization) during motor preparation was also associated with the consolidation of motor learning (Figure 6B). These findings provide novel evidence that brain network flexibility supports the establishment and stabilization of motor learning, although this contribution appears to be diminished with age, possibly reflecting age-related neural changes.

For the behavioral findings, YA and OA groups were able to adapt sufficiently to changes in the novel force mapping (i.e., previously unexperienced mapping). Although older adults had less precise cursor control compared to younger adults at the baseline (Supplementary Figure 1), they exhibited motor learning effects similar to those of younger adults as they could adapt to the novel mapping (from the baseline of 1:1 to the adaptation of 1:3) (Figure 2). As an overview of the motor learning task and definitions of the behavioral parameters, we considered the learning slope to reflect the magnitude of trial-by-trial changes in motor performance in response to experienced errors, whereas the aftereffect reflected the maintenance and recall of the internal model for future use. (Heuer and Hegele, 2008; Hegele and Heuer, 2010; Shadmehr et al., 2010; Krakauer and Mazzoni, 2011; Krakauer et al., 2019; Leech et al., 2022). From a neural perspective, previous studies suggest that neural activity emerging during the post-learning rest period is interpreted as the replay or reactivation of recently acquired motor memories (Buch et al., 2021; Griffin et al., 2025; Sjøgård et al., 2025). Accordingly, evaluating motor performance following a 10-min rest period may provide valuable insight into early offline processes that contribute to the stabilization and maintenance of internal models acquired from motor adaptation. We found that older adults exhibited a significantly steeper learning slope than younger adults (Figure 2B), whereas no age-related differences were observed in the aftereffect (Figure 2C). It can be interpreted as a scenario in which the rapid reduction in performance errors observed in older adults was unlikely to reflect an enhanced capacity for sensorimotor recalibration. Instead, it suggests that the steeper learning slope is primarily driven by larger trial-by-trial behavioral updates (Taylor et al., 2014; McDougle et al., 2015). This pattern may be explained by increased sensory noise from the periphery due to aging (Tran et al., 2020), which alters the weighting of prediction errors and results in larger trial-by-trial updates (van Beers et al., 2004). When faced with large visuomotor perturbations, older adults may increasingly recruit cognitive resources to deliberately adjust their aiming direction and speed control for the cursor on a trial-by-trial basis to counteract perceived errors (Cisneros et al., 2024). Building on these notable behavioral findings, we extend our discussion to the neurophysiological coupling underlying age-related motor learning.

Here, our findings highlight a neurophysiological link between brain network flexibility and learning characteristics, and this relationship was further modulated by age. Brain network flexibility during motor preparation and its reduction toward the later adaptation period showed the identical patterns associated with aging (Figure 6). In young adults, a progressive reduction in brain network flexibility toward later adaptation (i.e., network stabilization) was selectively associated with the stronger aftereffect, but not with learning rate. This indicates that network stabilization primarily supports post-learning retention of the adapted internal model. Indeed, this finding is consistent with previous evidence showing that better visual perception abilities are associated with larger quenching of neural variability (Daniel and Dinstein, 2021). In contrast, older adults showed the opposite pattern. The learning-related reductions in network flexibility were strongly associated with learning rate but were unrelated to the aftereffect. This dissociation suggests an age-related shift in the functional role of brain network flexibility from a post-learning stabilization mechanism in young adults to an online constraint on learning efficiency in older adults. This finding aligns with a previous human EEG study showing that older adults with larger trial-by-trial auditory response variability are less accurate at sound localization (Anderson et al., 2012), indicating that aging alters the stability of brain activity and these changes influence behavior. By contrast, such a strong association was not observed during motor execution in the visuomotor adaptation task (Figure 7). Accordingly, we suggest that large-scale network reconfiguration across the brain primarily supports planning and post-learning stabilization. In contrast, motor execution-related neural dynamics reflect the implementation of already specified or planned motor commands. This distinction arises because preparatory neural activity represents an early stage in the chain of events that generates voluntary movement (Churchland and Shenoy, 2024; Schimel et al., 2024). Contrary to our initial hypothesis, group-average brain network flexibility during motor preparation and execution did not differ between younger and older adults (Figure 4). However, age-related differences emerged in how brain network flexibility predicted individual differences in motor learning (Figures 6 and 7). This dissociation implies that aging may alter the functional significance of network flexibility at the individual level rather than its overall magnitude. Such an interpretation is broadly consistent with compensation-based theories of aging, including HAROLD, PASA, CRUNCH, and STAC (Cabeza, 2002; Davis et al., 2008; Reuter-Lorenz and Cappell, 2008; Reuter-Lorenz and Park, 2014; Son et al., 2024). These conceptual frameworks suggest that older adults maintain behavioral performance through the recruitment of alternative neural resources and reorganization of functional networks (Veldman et al., 2021), and aging affects the efficiency of brain networks for behavior (Lin et al., 2016; Ezaki et al., 2018; Battaglia et al., 2020; Deery et al., 2023). Under this framework, comparable levels of network flexibility may reflect distinct neural strategies across age groups, resulting in different neurobehavioral associations with motor learning. As an illustrative example, Yotsumoto et al. (Yotsumoto et al., 2014) reported a similar dissociation between neural activity and learning ability, indicating that perceptual learning relies on distinct neural mechanisms in older and younger adults, with older adults relying more on white matter reorganization than on functional brain changes. By combining previous evidence with our findings, motor learning in older adults may rely less on the brain network flexibility observed in younger adults and more on compensatory processes at other hierarchical levels of the nervous system.

We further discuss perspectives on the frequency-specific contributions of brain network flexibility to motor learning properties. Despite the complexity of interpreting frequency-specific effects, identifiable roles emerge from our findings. Our LASSO regression analyses revealed that the aftereffect was positively associated with alpha-band flexibility and negatively associated with theta- and beta-band flexibility in young adults. It is well established that alpha oscillations are ubiquitous in the brain (Berger, 1929), yet their functional significance remains incompletely understood. Recently, alpha oscillations are thought to play a central role in regulating network information flow (Sadaghiani et al., 2012; Scheeringa et al., 2012) and attention (Romei et al., 2012; Herring et al., 2015; Peylo et al., 2021). Rather than directly reflecting motor output, alpha-band dynamics have been linked to the establishment of task-ready network configurations and the gating of neural resources (Klimesch et al., 2007; Ai and Ro, 2014; Babiloni et al., 2014; Brickwedde et al., 2019; Borra et al., 2023). Therefore, the reduction in alpha-band flexibility observed in this study may reflect the stabilization of preparatory network states as learning progresses, reducing the need for continual large-scale network reconfiguration. Consistent with this interpretation, our predictive models using LASSO identified alpha-band brain network flexibility during motor preparation as a key predictor of the learning aftereffect (Figure 6).

As a final point, interpreting LZC as an index of neural complexity offers additional insight into our findings. We examined LZC because understanding local signal complexity is necessary for a more comprehensive understanding of brain network flexibility. The present study found a dissociation between resting-state LZC and brain network flexibility across the age groups, with higher LZC in older adults but comparable brain network flexibility (Figure 3). Drawing on previous evidence and our findings, we postulate that LZC is likely a sensitive indicator of local neural dynamics (Schartner et al., 2015; Medel et al., 2023) and clearly reflects the effects of aging (Fernández et al., 2012; Biggs et al., 2022). On the other hand, brain network flexibility may saturate owing to constraints on network reconfiguration, possibly because resilience mechanisms maintain large-scale network organization throughout the lifespan (Heisz et al., 2015; Gonzalez-Escamilla et al., 2018; Neudorf et al., 2024). This resilience mechanism and sensitivity to aging may lead to the resting-state dissociation. Another noteworthy finding is that the age-related elevations in LZC and preserved brain network flexibility were confined to the resting state and motor preparation, disappearing during motor execution. This selective manifestation suggests that the aging brain employs a proactive neural strategy characterized by hyper-complex idling. By maintaining a broader repertoire of dynamic states and high network reconfigurability before movement, older adults may create a functional buffer that compensates for age-related neurobiological declines (Goble et al., 2009; Coxon et al., 2014; Serbruyns et al., 2015; Frolov et al., 2020; Zapparoli et al., 2022; Goelman et al., 2023). This heightened readiness may facilitate the successful transition into a stabilized, task-optimal state during motor action, where motor performance becomes indistinguishable from that of the younger adults. Moreover, we statistically confirmed the significant positive correlations between LZC and brain network flexibility, as summarized in Table 1. This implies that both measures may be governed by a shared dynamical mechanism spanning local and global neural dynamics. It should be noted that the observed reductions in LZC for alpha- and beta-frequency bands might be partially attributed to the well-established event-related desynchronization (ERD) observed during motor behavior (Pfurtscheller and Lopes Da Silva, 1999). While ERD is conventionally characterized as a reduction in oscillatory power, complexity measures quantify the temporal irregularity of neural activity. Therefore, although ERD-related power changes and complexity changes may be linked, they do not necessarily reflect the same underlying neural phenomenon. Rather, they likely provide complementary information about cortical dynamics.

The present study has potential limitations and caveats regarding the interpretation of the results. First, we could not mention the cerebellar function. It is accepted that the cerebellum plays a key role in the regulation of motor learning, including adaptation (Krakauer and Mazzoni, 2011; Tsay et al., 2024). However, non-invasive measurement of cerebellar electrophysiology is hindered by the limited spatial resolution of human scalp EEG recordings (Andersen et al., 2020). In principle, studies employing EEG to delineate functional networks do not account for the cerebellum, as it is not reliably measurable using this technique. That said, the cerebellum is connected to broader cortical regions, including the motor, frontal, parietal cortices, and temporal lobe (Bernard et al., 2012; Schulz et al., 2014). Although we could not directly capture cerebellar function, assessing the dynamics of cortical networks, which receive extensive inputs from and send outputs to the cerebellum, is essential for understanding motor learning. Second, the present study compared only momentary differences between the two specific age groups. To account for the entire lifespan, the next step should be to employ a cross-sectional study design that assesses age-related changes along a continuum from infancy to old age.

In conclusion, the present study revealed dissociated roles of brain network flexibility in motor learning across the different age groups. Brain network flexibility during motor preparation, rather than motor execution, was critical for maintaining new motor action in younger adults, but this coupling was attenuated in older adults aged approximately 60 years and above. Understanding brain network flexibility and its state could facilitate the development of novel tools for enhancing motor learning and mitigating age-related learning limitations.

## Acknowledgments

We thank Assistant Professor Akiko Nishio from NIPS for her assistance in recruiting participants. This work was supported by JSPS KAKENHI Grant Number JP19K20103 and JST Grant Number JPMJPF2502 to K.U. and the grant of the OML Project by the National Institutes of Natural Sciences (NINS program No, OML032401) to KK.

## Conflict interests

The authors declare no conflicts of interest associated with this paper.

## Data and code availability

The source data and code underlying all main figures and Supplementary figures have been deposited in the Open Science Framework (OSF) via the following link https://doi.org/10.17605/OSF.IO/3FC94

Other data, including large raw continuous EEG data and behavior data, are available upon request from the corresponding authors.

## Supplementary Materials

**Supplementary Figure 1:**
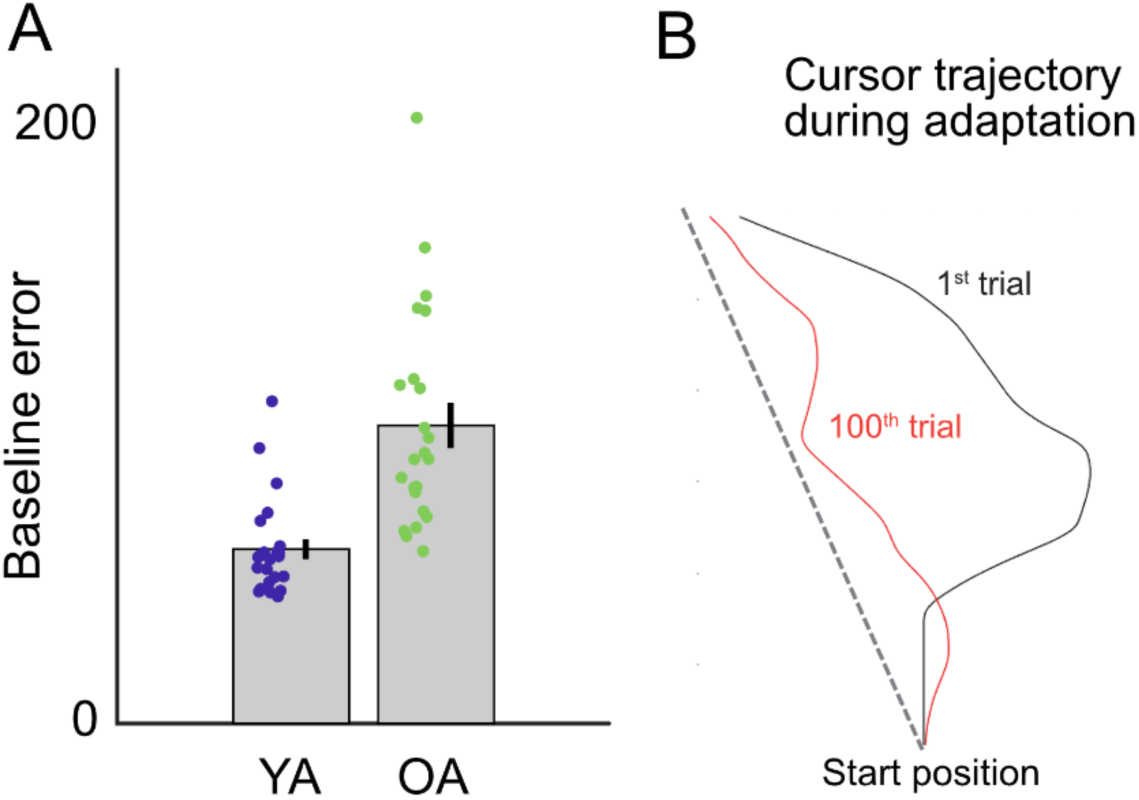
Baseline motor performance **(A)** and a typical cursor trajectory during adaptation **(B)**. **A**. Group-averaged error at the baseline across the fifty trials. Each dot represents data from an individual, and each error bar indicates the standard error of the mean across participants. **B**. A typical cursor trajectory during adaptation from a representative participant in the YA group. The deviation of the cursor was decreased consistently across the 100 trials.

## Notes

### Competing Interest Statement

The authors have declared no competing interest.

### Summary of Updates

We have fundamentally restructured the introduction and the discussion section. Specifically, we carefully restructured the discussion to ensure that our interpretations follow logically from the obtained findings.

## References

Ai L, Ro T (2014) The phase of prestimulus alpha oscillations affects tactile perception. J Neurophysiol 111:1300–1307.

Andersen LM, Jerbi K, Dalal SS (2020) Can EEG and MEG detect signals from the human cerebellum? Neuroimage 215:116817.

Anderson S, Parbery-Clark A, White-Schwoch T, Kraus N (2012) Aging affects neural precision of speech encoding. J Neurosci 32:14156–14164.

Arazi A, Censor N, Dinstein I (2017) Neural variability quenching predicts individual perceptual abilities. Journal of Neuroscience 37:97–109.

Babiloni C, Del Percio C, Arendt-Nielsen L, Soricelli A, Romani GL, Rossini PM, Capotosto P (2014) Cortical EEG alpha rhythms reflect task-specific somatosensory and motor interactions in humans. Clin Neurophysiol 125:1936–1945.

Bassett DS, Porter MA, Wymbs NF, Grafton ST, Carlson JM, Mucha PJ (2013) Robust detection of dynamic community structure in networks. Chaos 23:013142.

Bassett DS, Wymbs NF, Porter MA, Mucha PJ, Carlson JM, Grafton ST (2011) Dynamic reconfiguration of human brain networks during learning. Proc Natl Acad Sci U S A 108:7641–7646.

Battaglia D, Boudou T, Hansen ECA, Lombardo D, Chettouf S, Daffertshofer A, McIntosh AR, Zimmermann J, Ritter P, Jirsa V (2020) Dynamic Functional Connectivity between order and randomness and its evolution across the human adult lifespan. Neuroimage 222:117156.

Beck D, de Lange A-MG, Maximov II, Richard G, Andreassen OA, Nordvik JE, Westlye LT (2021) White matter microstructure across the adult lifespan: A mixed longitudinal and cross-sectional study using advanced diffusion models and brain-age prediction. Neuroimage 224:117441.

Berger H (1929) Über das elektroenkephalogramm des menschen. Arch Psychiatr Nervenkr 87:527–570.

Bernard J a., Seidler RD, Hassevoort KM, Benson BL, Welsh RC, Wiggins JL, Jaeggi SM, Buschkuehl M, Monk CS, Jonides J, Peltier SJ (2012) Resting state cortico-cerebellar functional connectivity networks: a comparison of anatomical and self-organizing map approaches. Front Neuroanat 6:31–31.

Betzel RF, Satterthwaite TD, Gold JI, Bassett DS (2017) Positive affect, surprise, and fatigue are correlates of network flexibility. Sci Rep 7:1–10.

Biggs D, Boncompte G, Pedemonte JC, Fuentes C, Cortinez LI (2022) The effect of age on electroencephalogram measures of anesthesia hypnosis: A comparison of BIS, Alpha Power, Lempel-Ziv complexity and permutation entropy during propofol induction. Front Aging Neurosci 14:910886.

Blondel VD, Guillaume J-L, Lambiotte R, Lefebvre E (2008) Fast unfolding of communities in large networks. J Stat Mech 2008:P10008.

Bönstrup M, Iturrate I, Hebart MN, Censor N, Cohen LG (2020) Mechanisms of offline motor learning at a microscale of seconds in large-scale crowdsourced data. npj Science of Learning 5:1–10.

Bönstrup M, Iturrate I, Thompson R, Cruciani G, Censor N, Cohen LG (2019) A rapid form of offline consolidation in skill learning. Curr Biol 29:1346–1351.e4.

Borra D, Fantozzi S, Bisi MC, Magosso E (2023) Modulations of cortical power and connectivity in alpha and beta bands during the preparation of reaching movements. Sensors (Basel) 23:3530.

Braun U, Schäfer A, Walter H, Erk S, Romanczuk-Seiferth N, Haddad L, Schweiger JI, Grimm O, Heinz A, Tost H, Meyer-Lindenberg A, Bassett DS (2015) Dynamic reconfiguration of frontal brain networks during executive cognition in humans. Proc Natl Acad Sci U S A 112:11678–11683.

Brickwedde M, Krüger MC, Dinse HR (2019) Somatosensory alpha oscillations gate perceptual learning efficiency. Nat Commun 10:263.

Buch ER, Claudino L, Quentin R, Bönstrup M, Cohen LG (2021) Consolidation of human skill linked to waking hippocampo-neocortical replay. Cell Rep 35:109193.

Cabeza R (2002) Hemispheric asymmetry reduction in older adults: the HAROLD model. Psychol Aging 17:85–100.

Churchland MM, Shenoy KV (2024) Preparatory activity and the expansive null-space. Nat Rev Neurosci 25:213–236.

Cisneros E, Karny S, Ivry RB, Tsay JS (2024) Differential aging effects on implicit and explicit sensorimotor learning. bioRxivorg Available at: 10.1101/2024.07.02.601091.

Cohen MX (2014) Analyzing neural time series data: Theory and practice. Mit Press.

Cohen MX (2019) A better way to define and describe Morlet wavelets for time-frequency analysis. Neuroimage 199:81–86.

Courchesne E, Chisum HJ, Townsend J, Cowles A, Covington J, Egaas B, Harwood M, Hinds S, Press GA (2000) Normal brain development and aging: quantitative analysis at in vivo MR imaging in healthy volunteers. Radiology 216:672–682.

Cox SR, Ritchie SJ, Tucker-Drob EM, Liewald DC, Hagenaars SP, Davies G, Wardlaw JM, Gale CR, Bastin ME, Deary IJ (2016) Ageing and brain white matter structure in 3,513 UK Biobank participants. Nat Commun 7:13629.

Coxon JP, Goble DJ, Leunissen I, Van Impe A, Wenderoth N, Swinnen SP (2014) Functional Brain Activation Associated with Inhibitory Control Deficits in Older Adults. Cereb Cortex:1–11.

Daniel E, Dinstein I (2021) Individual magnitudes of neural variability quenching are associated with motion perception abilities. J Neurophysiol 125:1111–1120.

Davis SW, Dennis NA, Daselaar SM, Fleck MS, Cabeza R (2008) Que PASA? The posterior-anterior shift in aging. Cereb Cortex 18:1201–1209.

Deery HA, Di Paolo R, Moran C, Egan GF, Jamadar SD (2023) The older adult brain is less modular, more integrated, and less efficient at rest: A systematic review of large-scale resting-state functional brain networks in aging. Psychophysiology 60:e14159.

Delorme A, Makeig S (2004) EEGLAB: An open source toolbox for analysis of single-trial EEG dynamics including independent component analysis. J Neurosci Methods 134:9–21.

Dudai Y (2004) The neurobiology of consolidations, or, how stable is the engram? Annu Rev Psychol 55:51–86.

Erdfelder E, FAul F, Buchner A, Lang AG (2009) Statistical power analyses using G*Power 3.1: Tests for correlation and regression analyses. Behav Res Methods 41:1149–1160.

Ezaki T, Sakaki M, Watanabe T, Masuda N (2018) Age-related changes in the ease of dynamical transitions in human brain activity. Hum Brain Mapp 39:2673–2688.

Fernández A, López-Ibor M-I, Turrero A, Santos J-M, Morón M-D, Hornero R, Gómez C, Méndez MA, Ortiz T, López-Ibor JJ (2011) Lempel-Ziv complexity in schizophrenia: a MEG study. Clin Neurophysiol 122:2227–2235.

Fernández A, Zuluaga P, Abásolo D, Gómez C, Serra A, Méndez MA, Hornero R (2012) Brain oscillatory complexity across the life span. Clin Neurophysiol 123:2154–2162.

Folstein MF, Folstein SE, McHugh PR (1975) “Mini-mental state”. A practical method for grading the cognitive state of patients for the clinician. J Psychiatr Res 12:189–198.

Friedman JH, Hastie T, Tibshirani R (2010) Regularization Paths for Generalized Linear Models via Coordinate Descent. J Stat Softw 33:1–22.

Frolov NS, Pitsik EN, Maksimenko VA, Grubov VV, Kiselev AR, Wang Z, Hramov AE (2020) Age-related slowing down in the motor initiation in elderly adults. PLoS One 15:e0233942.

Garrett DD, Kovacevic N, McIntosh AR, Grady CL (2010) Blood Oxygen Level-Dependent Signal Variability Is More than Just Noise. Journal of Neuroscience 30:4914–4921.

Garrett DD, Kovacevic N, McIntosh AR, Grady CL (2013) The modulation of BOLD variability between cognitive states varies by age and processing speed. Cereb Cortex 23:684–693.

Goble DJ, Coxon JP, Wenderoth N, Van Impe A, Swinnen SP (2009) Proprioceptive sensibility in the elderly: degeneration, functional consequences and plastic-adaptive processes. Neurosci Biobehav Rev 33:271–278.

Goelman G, Dan R, Bezdicek O, Jech R (2023) Directed functional connectivity of the sensorimotor system in young and older individuals. Front Aging Neurosci 15:1222352.

Gonzalez-Escamilla G, Muthuraman M, Chirumamilla VC, Vogt J, Groppa S (2018) Brain networks reorganization during maturation and healthy aging-emphases for resilience. Front Psychiatry 9:601.

Good CD, Johnsrude IS, Ashburner J, Henson RN, Friston KJ, Frackowiak RS (2001) A voxel-based morphometric study of ageing in 465 normal adult human brains. Neuroimage 14:21–36.

Griffin S, Khanna P, Choi H, Thiesen K, Novik L, Morecraft RJ, Ganguly K (2025) Ensemble reactivations during brief rest drive fast learning of sequences. Nature Available at: 10.1038/s41586-024-08414-9.

Guitart-Masip M, Salami A, Garrett D, Rieckmann A, Lindenberger U, Bäckman L (2016) BOLD Variability is Related to Dopaminergic Neurotransmission and Cognitive Aging. Cereb Cortex 26:2074–2083.

Haith AM, Huberdeau DM, Krakauer JW (2015) The influence of movement preparation time on the expression of visuomotor learning and savings. J Neurosci 35:5109–5117.

Hegele M, Heuer H (2010) The impact of augmented information on visuo-motor adaptation in younger and older adults. PLoS One 5:e12071.

Hehl M, Swinnen SP, Cuypers K (2020) Alterations of hand sensorimotor function and cortical motor representations over the adult lifespan. Aging (Albany NY) 12:4617–4640.

Heisz JJ, Gould M, McIntosh AR (2015) Age-related shift in neural complexity related to task performance and physical activity. J Cogn Neurosci 27:605–613.

Hermans L, Levin O, Maes C, van Ruitenbeek P, Heise KF, Edden RAE, Puts NAJ, Peeters R, King BR, Meesen RLJ, Leunissen I, Swinnen SP, Cuypers K (2018) GABA levels and measures of intracortical and interhemispheric excitability in healthy young and older adults: an MRS-TMS study. Neurobiol Aging 65:168–177.

Herring JD, Thut G, Jensen O, Bergmann TO (2015) Attention Modulates TMS-Locked Alpha Oscillations in the Visual Cortex. J Neurosci 35:14435–14447.

Heuer H, Hegele M (2008) Constraints on visuo-motor adaptation depend on the type of visual feedback during practice. Exp Brain Res 185:101–110.

Holland BS, Di Ponzio Copenhaver M (1988) Improved Bonferroni-Type Multiple Testing Procedures. Psychological Bulletin 104:145–149.

Jernigan TL, Archibald SL, Fennema-Notestine C, Gamst AC, Stout JC, Bonner J, Hesselink JR (2001) Effects of age on tissues and regions of the cerebrum and cerebellum. Neurobiol Aging 22:581–594.

Kaasinen V, Rinne JO (2002) Functional imaging studies of dopamine system and cognition in normal aging and Parkinson’s disease. Neurosci Biobehav Rev 26:785–793.

Kayser J, Tenke CE (2006) Principal components analysis of Laplacian waveforms as a generic method for identifying ERP generator patterns: II. Adequacy of low-density estimates. Clin Neurophysiol 117:369–380.

Klimesch W, Sauseng P, Hanslmayr S (2007) EEG alpha oscillations: the inhibition-timing hypothesis. Brain Res Rev 53:63–88.

Krakauer JW, Hadjiosif AM, Xu J, Wong AL, Haith AM (2019) Motor Learning. Compr Physiol 9:613–663.

Krakauer JW, Mazzoni P (2011) Human sensorimotor learning: Adaptation, skill, and beyond. Curr Opin Neurobiol 21:636–644.

Krukow P, Rodríguez-González V, Kopiś-Posiej N, Gómez C, Poza J (2024) Tracking EEG network dynamics through transitions between eyes-closed, eyes-open, and task states. Sci Rep 14:17442.

Lachaux J-P, Rodriguez E, M LVQ, A L, J M, Varela FJ (2000) Studying signale-trials of phase synchronous activity in the brain. Int J Bifurcat Chaos 10:2429–2439.

Lachaux JP, Rodriguez E, Martinerie J, Varela FJ (1999) Measuring phase synchrony in brain signals. Hum Brain Mapp 8:194–208.

Leech KA, Roemmich RT, Gordon J, Reisman DS, Cherry-Allen KM (2022) Updates in Motor Learning: Implications for Physical Therapist Practice and Education. Phys Ther 102 Available at: 10.1093/ptj/pzab250.

Lempel A, Ziv J (1976) On the complexity of finite sequences. IEEE Trans Inf Theory 22:75–81.

Leow L-A, Marinovic W, de Rugy A, Carroll TJ (2018) Task errors contribute to implicit aftereffects in sensorimotor adaptation. Eur J Neurosci 48:3397–3409.

Lin C-HJ, Knowlton BJ, Wu AD, Iacoboni M, Yang H-C, Ye Y-L, Liu K-H, Chiang M-C (2016) Benefit of interleaved practice of motor skills is associated with changes in functional brain network topology that differ between younger and older adults. Neurobiol Aging 42:189–198.

Liuzzi L, Quinn AJ, O’Neill GC, Woolrich MW, Brookes MJ, Hillebrand A, Tewarie P (2019) How sensitive are conventional MEG functional connectivity metrics with sliding windows to detect genuine fluctuations in dynamic functional connectivity? Front Neurosci 13:797.

Marstaller L, Williams M, Rich A, Savage G, Burianová H (2015) Aging and large-scale functional networks: white matter integrity, gray matter volume, and functional connectivity in the resting state. Neuroscience 290:369–378.

McDougle SD, Bond KM, Taylor JA (2015) Explicit and implicit processes constitute the fast and slow processes of sensorimotor learning. J Neurosci 35:9568–9579.

Medel V, Irani M, Crossley N, Ossandón T, Boncompte G (2023) Complexity and 1/f slope jointly reflect brain states. Sci Rep 13:21700.

Mognon A, Jovicich J, Bruzzone L, Buiatti M (2011) ADJUST: An automatic EEG artifact detector based on the joint use of spatial and temporal features. Psychophysiology 48:229–240.

Monteiro TS, King BR, Zivari Adab H, Mantini D, Swinnen SP (2019) Age-related differences in network flexibility and segregation at rest and during motor performance. Neuroimage 194:93–104.

Mucha PJ, Richardson T, Macon K, Porter MA, Onnela J-P (2010) Community structure in time-dependent, multiscale, and multiplex networks. Science 328:876–878.

Neudorf J, Shen K, McIntosh AR (2024) Reorganization of structural connectivity in the brain supports preservation of cognitive ability in healthy aging. Netw Neurosci 8:837–859.

Oldfield RC (1971) The assessment and analysis of handedness: The Edinburgh inventory. Neuropsychologia 9:97–113.

O’Sullivan I, Burdet E, Diedrichsen J (2009) Dissociating Variability and Effort as Determinants of Coordination. PLoS Comput Biol 5 Available at: 10.1371/journal.pcbi.1000345.

Paban V, Modolo J, Mheich A, Hassan M (2019) Psychological resilience correlates with EEG source-space brain network flexibility. Netw Neurosci 3:539–550.

Parra D, Zhang Z, Radvansky G (2026) Should we all just take 10? A meta-analysis of wakeful rest. Psychon Bull Rev 33:49.

Pauelsen M, Jafari H, Strandkvist V, Nyberg L, Gustafsson T, Vikman I, Röijezon U (2020) Frequency domain shows: Fall-related concerns and sensorimotor decline explain inability to adjust postural control strategy in older adults. PLoS One 15:e0242608.

Perrin F, Pernier J, Bertrand O, Echallier JF (1989) Spherical splines for scalp potential and current density mapping. Electroencephalogr Clin Neurophysiol 72:184–187.

Peylo C, Hilla Y, Sauseng P (2021) Cause or consequence? Alpha oscillations in visuospatial attention. Trends Neurosci 44:705–713.

Pfurtscheller G, Lopes Da Silva FH (1999) Event-related EEG/MEG synchronization and desynchronization: Basic principles. Clin Neurophysiol 110:1842–1857.

Ranganathan R, Lee M-H, Newell KM (2020) Repetition Without Repetition: Challenges in Understanding Behavioral Flexibility in Motor Skill. Front Psychol 11:2018.

Ranganathan R, Newell KM (2010) Emergent flexibility in motor learning. Exp Brain Res 202:755–764.

Reddy PG, Mattar MG, Murphy AC, Wymbs NF, Grafton ST, Satterthwaite TD, Bassett DS (2018) Brain state flexibility accompanies motor-skill acquisition. Neuroimage 171:135–147.

Reuter-Lorenz PA, Cappell KA (2008) Neurocognitive aging and the compensation hypothesis. Curr Dir Psychol Sci 17:177–182.

Reuter-Lorenz PA, Park DC (2014) How does it STAC up? Revisiting the scaffolding theory of aging and cognition. Neuropsychol Rev 24:355–370.

Rodriguez E, George N, Lachaux JP, Martinerie J, Renault B, Varela FJ (1999) Perception’s shadow: Long-distance synchronization of human brain activity. Nature 397:430–433.

Romei V, Thut G, Mok RM, Schyns PG, Driver J (2012) Causal implication by rhythmic transcranial magnetic stimulation of alpha frequency in feature-based local vs. global attention. Eur J Neurosci 35:968–974.

Sadaghiani S, Scheeringa R, Lehongre K, Morillon B, Giraud A-L, D’Esposito M, Kleinschmidt A (2012) Alpha-Band Phase Synchrony Is Related to Activity in the Fronto-Parietal Adaptive Control Network. Journal of Neuroscience 32:14305–14310.

Salomonczyk D, Cressman EK, Henriques DYP (2013) The role of the cross-sensory error signal in visuomotor adaptation. Exp Brain Res 228:313–325.

Schartner M, Seth A, Noirhomme Q, Boly M, Bruno M-A, Laureys S, Barrett A (2015) Complexity of multi-dimensional spontaneous EEG decreases during propofol induced general anaesthesia. PLoS One 10:e0133532.

Scheeringa R, Petersson KM, Kleinschmidt A, Jensen O, Bastiaansen MCM (2012) EEG Alpha Power Modulation of fMRI Resting-State Connectivity. Brain Connect 2:254–264.

Schimel M, Kao T-C, Hennequin G (2024) When and why does motor preparation arise in recurrent neural network models of motor control? Elife 12 Available at: 10.7554/eLife.89131.

Schulz R, Wessel MJ, Zimerman M, Timmermann JE, Gerloff C, Hummel FC (2014) White Matter Integrity of Specific Dentato-Thalamo-Cortical Pathways is Associated with Learning Gains in Precise Movement Timing. Cereb Cortex:1–8.

Serbruyns L, Gooijers J, Caeyenberghs K, Meesen RL, Cuypers K, Sisti HM, Leemans A, Swinnen SP (2015) Bimanual motor deficits in older adults predicted by diffusion tensor imaging metrics of corpus callosum subregions. Brain Struct Funct:1–18.

Shadmehr R, Smith MA, Krakauer JW (2010) Error correction, sensory prediction, and adaptation in motor control. Annu Rev Neurosci 33:89–108.

Shafiei G, Zeighami Y, Clark CA, Coull JT, Nagano-Saito A, Leyton M, Dagher A, Mišic B (2019) Dopamine signaling modulates the stability and integration of intrinsic brain networks. Cereb Cortex 29:397–409.

Sjøgård M, Baxter B, Mylonas D, Thompson M, Kwok K, Driscoll B, Tolosa A, Shi W, Stickgold R, Vangel M, Chu CJ, Manoach DS (2025) Hippocampal ripples predict motor learning during brief rest breaks in humans. Nat Commun 16:6089.

Son JJ, Arif Y, Okelberry HJ, Johnson HJ, Willett MP, Wiesman AI, Wilson TW (2024) Aging modulates the impact of cognitive interference subtypes on dynamic connectivity across a distributed motor network. NPJ Aging 10:54.

Sun X, O’Shea DJ, Golub MD, Trautmann EM, Vyas S, Ryu SI, Shenoy KV (2022) Cortical preparatory activity indexes learned motor memories. Nature:1–6.

Tallon-Baudry C, Bertrand O, Delpuech C, Pernier J (1996) Stimulus specificity of phase-locked and non-phase-locked 40 Hz visual responses in human. J Neurosci 16:4240–4249.

Taylor JA, Krakauer JW, Ivry RB (2014) Explicit and implicit contributions to learning in a sensorimotor adaptation task. J Neurosci 34:3023–3032.

Tibshirani R (1996) Regression shrinkage and selection via the lasso. J R Stat Soc 58:267–288.

Tran TT, Rolle CE, Gazzaley A, Voytek B (2020) Linked sources of neural noise contribute to age-related cognitive decline. J Cogn Neurosci 32:1813–1822.

Tsay JS, Kim HE, McDougle SD, Taylor JA, Haith A, Avraham G, Krakauer JW, Collins AGE, Ivry RB (2024) Fundamental processes in sensorimotor learning: Reasoning, refinement, and retrieval. Elife 13 Available at: 10.7554/eLife.91839.

Uehara K, Yasuhara M, Koguchi J, Oku T, Shiotani S, Morise M, Furuya S (2023) Brain network flexibility as a predictor of skilled musical performance. Cereb Cortex Available at: 10.1093/cercor/bhad298.

Uehara S, Mawase F, Therrien AS, Cherry-Allen KM, Celnik P (2019) Interactions between motor exploration and reinforcement learning. J Neurophysiol 122:797–808.

van Beers RJ, Haggard P, Wolpert DM (2004) The role of execution noise in movement variability. J Neurophysiol 91:1050–1063.

van de Vijver I, Cohen MX (2019) Electrophysiological phase synchrony in distributed brain networks as a promising tool in the study of cognition. In: New Methods in Cognitive Psychology (Spieler D, Schumacher E, eds), pp 214–244 1st Edition. Routledge.

van Diessen E, Numan T, van Dellen E, van der Kooi AW, Boersma M, Hofman D, van Lutterveld R, van Dijk BW, van Straaten ECW, Hillebrand A, Stam CJ (2015) Opportunities and methodological challenges in EEG and MEG resting state functional brain network research. Clin Neurophysiol 126:1468–1481.

Varela F, Lachaux JP, Rodriguez E, Martinerie J (2001) The brainweb: phase synchronization and large-scale integration. Nat Rev Neurosci 2:229–239.

Veldman MP, Maurits NM, Mantini D, Hortobágyi T (2021) Age-dependent modulation of motor network connectivity for skill acquisition, consolidation and interlimb transfer after motor practice. Clin Neurophysiol 132:1790–1801.

Vercillo T, Carrasco C, Jiang F (2017) Age-related changes in sensorimotor temporal binding. Front Hum Neurosci 11:500.

Wu HG, Miyamoto YR, Nicolas L, Castro G, Ölveczky BP, Smith MA, Biology E (2014) Temporal structure of motor variability is dynamically regulated and predicts motor learning ability. Nat Neurosci 17:312–321.

Yoshimura N, Tsuda H, Aquino D, Takagi A, Ogata Y, Koike Y, Minati L (2020) Age-related decline of sensorimotor integration influences resting-state functional brain connectivity. Brain Sci 10:966.

Yotsumoto Y, Chang L-H, Ni R, Pierce R, Andersen GJ, Watanabe T, Sasaki Y (2014) White matter in the older brain is more plastic than in the younger brain. Nat Commun 5:5504.

Zapparoli L, Mariano M, Paulesu E (2022) How the motor system copes with aging: a quantitative meta-analysis of the effect of aging on motor function control. Commun Biol 5:79.

